# The protein translation initiation machinery is targetable for developing fast-killing therapeutics against the zoonotic *Cryptosporidium parvum*

**DOI:** 10.1101/2025.01.03.631182

**Authors:** Meng Li, Jigang Yin, Dongqiang Wang, Beibei Zou, Pingwei Li, Guan Zhu

## Abstract

*Cryptosporidium parvum* is a zoonotic apicomplexan parasite that causes moderate to severe watery diarrhea in humans and animals. It is one of the two major cryptosporidial species that infect humans, particularly those with weak or compromised immunity (e.g., children and AIDS patients). *C. parvum* is also one of the major pathogens causing diarrhea in neonatal farm ruminants (e.g., calves). However, fully effective drugs are still unavailable for medical or veterinary use to treat cryptosporidiosis. In the past decade or two, a number of key enzymes in the core metabolic pathways of *Cryptosporidium* have been explored as drug targets (e.g., energy metabolism, fatty acid synthesis, nucleotide synthesis, protein synthesis and modification), while the protein translation initiation machinery has been unexplored. Here, we report that *C*. *parvum* eukaryotic initiation factor 4A (CpeIF4A), a DEAD-box RNA helicase and part of the eIF4F complex (composed of eIF4A, eIF4E, and eIF4G), can be targeted for developing anti-cryptosporidial therapeutics. CpeIF4A can be inhibited by the selective inhibitor rocaglamide-A (Roc-A; a natural product from *Aglaia* plants). The inhibition of CpeIF4A activity by Roc-A impedes protein synthesis in the sporozoites of *C. parvum* (IC_50_ = 3.65–3.80 μM) via tight binding to the CpeIF4A–RNA complex (K_d_ = 33.7 nM). This efficaciously lowers parasite growth in vitro (EC_50_ = 1.77 nM; selectivity index >1000) and oocyst production in vivo in IFN-γ knockout mice (with a rapid drop on the second day post-administration at 0.5 mg/kg/day).

These findings indicate that CpeIF4A can serve as a novel drug target, and that Roc-A is a promising anti-cryptosporidial lead compound for therapeutic development and structural optimization.

**Author summary:** *Cryptosporidium parvum* is a globally distributed protozoan parasite that infect both humans and animals. Cryptosporidial infection in hosts with weak or compromised immunity (e.g., AIDS patients, children or calves) can be severe or deadly, but fully effective treatments are yet unavailable. In the past decade or two, a number of potential drug targets in the core metabolic pathways in *Cryptosporidium* have been explored, but the protein translation machinery has been unexplored. Here, we report that the *C. parvum* eukaryotic initiation factor 4A (CpeIF4A), one of the key components in the protein translational machinery, can be targeted for developing fast-killing anti-cryptosporidial therapeutics. We show that CpeIF4A can be inhibited by the selective inhibitor rocaglamide-A (Roc-A), consequently suppressing the protein synthesis in the parasite via tight binding to the CpeIF4A–RNA complex. This efficaciously inhibits the parasite growth both in vitro and in vivo. The killing of *C. parvum* by Roc-A is fast and irreversible (parasiticidal). Our data indicate that the protein translational machinery is a novel drug target for developing fast-killing anti-cryptosporidial therapeutics, and that Roc-A is a promising lead compound for therapeutic development and structural optimization.

## Introduction

*Cryptosporidium* is a genus of widely distributed enteric apicomplexan parasites. Of more than 40 named *Cryptosporidium* species, *C. parvum* and *C. hominis* are the most medically important, responsible for the majority of the human cases of cryptosporidiosis [1]. Cryptosporidial infection is typically acute and self-limiting in immunocompetent patients, by which the diarrheal and associated symptoms may last a week or two; but can be prolonged and severe, and sometime deadly, in people in more vulnerable population, including those with weakened or compromised immunity [2,3]. In resource-restricted countries, cryptosporidiosis is one of the top diarrheal-causing agents in children and associated with stunt growth and increased death rate [3–5]. The zoonotic *C. parvum* also infects other mammals, particularly newborn ruminants (e.g., calves), which not only cause significant economic loss in the cattle industry [6,7], but also serve as a reservoir for human infection [8,9]. However, current options to treat cryptosporidiosis in humans and animals remain limited [10,11]. Nitazoxanide is the only drug approved by the United States Federal Drug Administration (FDA) to treat cryptosporidiosis in immunocompetent patients. Moreover, nitazoxanide is not fully effective or ineffective in immunocompromised individuals. Therefore, there is an urgent need to develop new anti-cryptosporidial drugs, particularly those for use in children and immunocompromised patients, and for veterinary use as well [11,12].

The slow progress in anti-cryptosporidial drug discovery is largely attributed to some technical difficulties and unique parasite biology [10]. Technically, *Cryptosporidium* cannot complete life cycle under routine in vitro conditions. Genetic manipulation tools are available, but still inconvenient for routine operation [13,14]. Biologically, *Cryptosporidium* differs from other apicomplexans (e.g., *Eimeria*, *Cyclospora*, *Toxoplasma* and *Plasmodium*) by lacking an apicoplast and associated metabolic pathways, the majority of the mitochondrial function (e.g., absence of Krebs cycle and respiratory chain in the intestinal *Cryptosporidium* species), and the pathways to synthesize any nutrients de novo (e.g., nucleosides, amino acids and fatty acids) [15,16]. These result in the absence of many traditional drug targets that are present in the coccidia and hematozoa lineages. Additionally, *Cryptosporidium* resides on top of the enterocytes, contained in host cell-derived parasitophorous vacuole membrane (PVM), but separated from the host cell by an electron-dense band and a membrane structure called feeder organelle (FO) [17,18]. These structures at the parasite-host cell interface are selective in permeability, which may block the access of certain small molecules to the parasite.

In target-based drug discovery, increased efforts have been devoted to discovery of selective inhibitors that target the key enzymes and factors in the core metabolic pathways present in *Cryptosporidium* in the past decade or two, such as those in the energy metabolism, fatty acid activation and elongation, nucleotide synthesis, protein synthesis, and protein modification [19–23]. The efforts lead to the discovery of many anti-cryptosporidial hits or leads, particularly those with defined inhibitory kinetics on the targets, excellent in vitro efficacy and selectivity profiles, and satisfactory in vivo efficacy [11,24]. Example leads include triacsin C that acts on *C. parvum* fatty acyl-CoA synthetase (ACS) [25], vorinostat on histone deacetylase (HDAC) [26], bumped kinase inhibitors (e.g., BKI-1708 and BKI-1553) on calcium-dependent protein kinase (CDPK) [27], KDU731 on phosphatidylinositol 4-kinase (PI4K), and compound 5 on Lysyl-tRNA synthetase (KRS) [23]. These targets function in core metabolism and biological processes essential to the parasite growth and development.

Here, we report that the protein translational initiation machinery in *Cryptosporidium*, which was previously unexplored, can be targeted for developing anti-cryptosporidial therapeutics. More specifically, we have found that the *C. parvum* eukaryotic initiation factor 4A (CpeIF4A), a DEAD-box (DDX) RNA helicase and part of the eIF4F complex comprised of eIF4A, eIF4E, and eIF4G, can be inhibited by the selective inhibitor rocaglamide-A (Roc-A; a natural product from *Aglaia* plants) (**Fig. 1A**) [28]. The inhibition of CpeIF4A activity by Roc-A impedes the protein synthesis in the sporozoites of *C. parvum*, efficaciously lowering the parasite growth in vitro and the oocyst production in interferon-γ-knockout (IFN-γ KO) mice. These findings indicate that CpeIF4A can serve as a novel drug target, and Roc-A is worth being pursued as a new anti-cryptosporidial lead for therapeutical development and/or structural optimization.

**Fig. 1.**
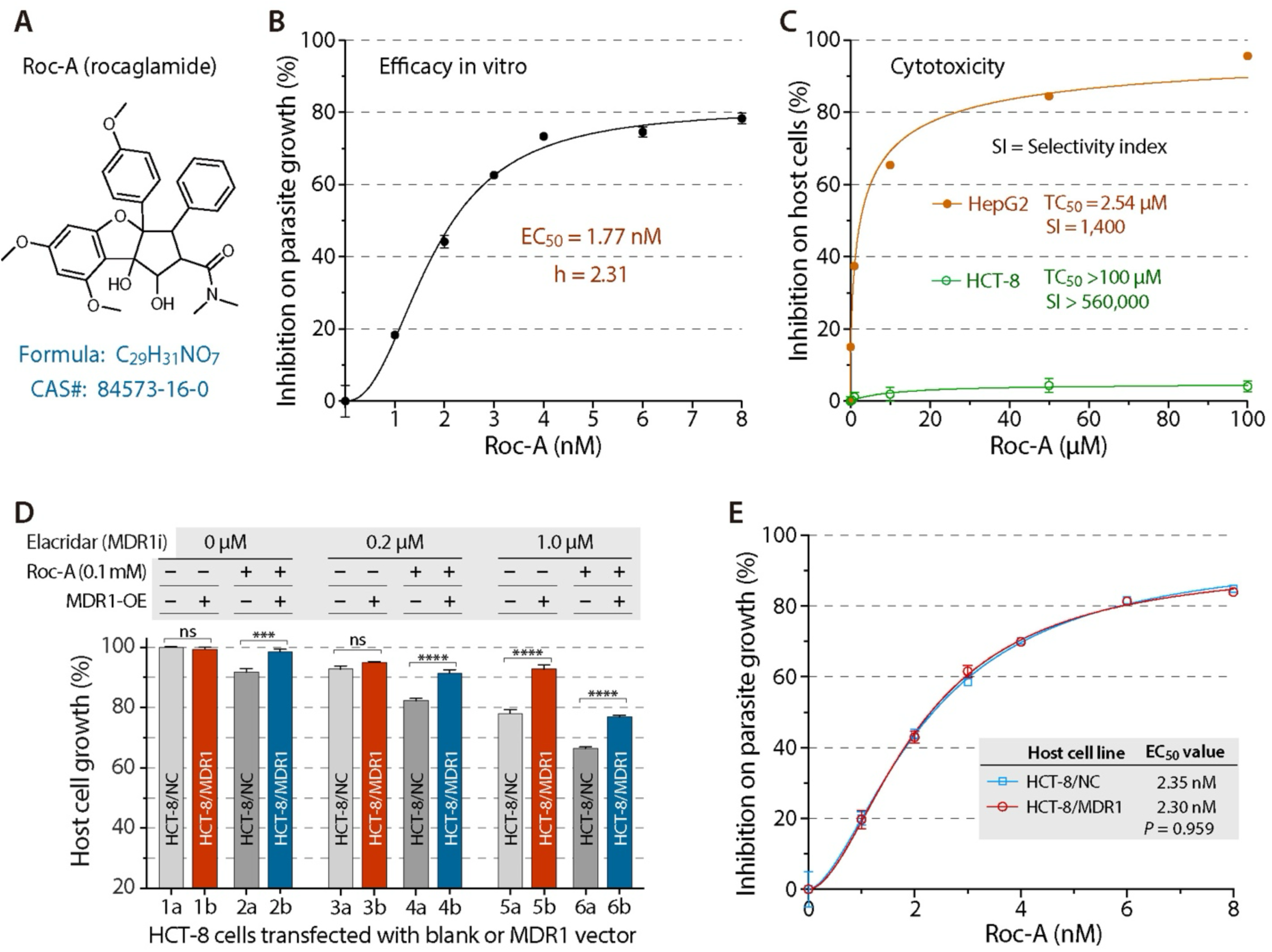
Selective and on-the-parasite-target inhibition of rocaglamide (Roc-A) against the growth of *C. parvum* in vitro. **(A)** Chemical structure of Roc-A (CAS# 84573-16-0), a highly selective inhibitor of eukaryotic initiation factor 4A (eIF4A). **(B)** In vitro efficacy of Roc-A against the growth of *C. parvum* grown on HCT-8 cells in a 44-h infection assay. The parasite loads were quantified by qRT-PCR that detects the *C. parvum* 18S rRNA (Cp18S) levels. **(C)** Cytotoxicity of Roc-A on HCT-8 (host cells) and hepG2 cells. Cell proliferation was assessed after 44-h treatment by MTS assay. Selectivity index (SI) indicates the ratio between TC_50_ (50% toxic concentration on HCT-8 or HepG2 cells) and EC_50_ (50% effective inhibition concentration on *C. parvum*) (i.e., SI = TC_50_/EC_50_). **(D)** Susceptibility of HCT-8 cells transiently transfected with human *MDR1* gene (marked as HCT-8/MDR1) in comparison to the blank vector negative control (HCT-8/NC) to the treatment of Roc-A (0.1 mM) in the absence or presence of elacridar (a third generation inhibitor of MDR1). In this assay, MDR1 over-expression (MDR1-OE) increases the tolerance of HCT-8 cells to Roc-A in all elacridar treatment groups (HCT-8/MDR1 vs. HCT-8/NC). **(E)** Effect of MDR1 over-expression on the efficacy of Roc-A against *C. parvum*. The increase of drug tolerance of HCT-8 cells by the over-expression of MDR1 has no effect on the inhibitory curves and EC_50_ values (HCT-8/MDR1 vs. HCT-8/NC), indicating the inhibition of Roc-A on *C. parvum* is attributed to the on-the-parasite target effect.

## Results

### The eIF4A inhibitor Roc-A is highly efficacious and selective against the in vitro growth of *C. parvum* with confirmed effect on the parasite target

To test whether the translation initiation machinery could serve as a rational drug target in *C. parvum*, we first tested the anti-cryptosporidial efficacy of Roc-A in vitro, which is known for specific inhibition of eIF4A, a factor required for the binding of the capped mRNA to 40S ribosomal subunits to initiate the protein translation process [28]. Using an in vitro model, in which *C. parvum* was cultured in HCT-8 cells and the parasite loads were evaluated by qRT-PCR at 44 h post-infection (hpi) timepoint [29,30], Roc-A displays exceptionally high potence in inhibiting the parasite growth, showing lower single-digit nanomolar anti-parasitic activity (i.e., the 50% effective concentration (EC_50_ = 1.77 nM) (**Fig. 1B**). In contrast, the host cells (HCT-8 cell line derived from a human ileocecal colorectal adenocarcinoma) were much more tolerant to Roc-A as determined by MTS assay (**Fig. 1C**), in which little cytotoxicity was observed at concentrations up to 100 μM (i.e., the 50% toxic concentration (TC_50_) is much higher than 100 μM). This gives an exceptionally high selectivity index (SI) (i.e., SI is much higher than 560,000). The hepatocellular carcinoma HapG2 cells is less tolerant to Roc-A than HCT-8 cells, but still shows very high, 1,400-fold selectivity (i.e., TC_50_ = 2.54 μM; SI = 1,400).

The extremely high selectivity is indicative that the anti-cryptosporidial efficacy is fully attributed to the direct action of Roc-A on the parasite, rather than indirectly via acting on the host cells [31]. This notion is further confirmed using a recently developed in vitro model, in which HCT-8 cells are transiently or stably transfected to overexpress the multi-drug resistance-1 gene (*MDR1*) (HCT-8/MDR1; vs. the negative control carrying blank vector, or HCT-8/NC). In this model, if a compound is a substrate of MDR1, host cells overexpressing MDR1 will increase drug tolerance to the compound. If the compound acts fully on the epicellular *Cryptosporidium* (i.e., on the parasite target), the increased drug tolerance in host cells would not alter the anti-cryptosporidial efficacy of the compound [31,32]. Here, transient overexpression of MDR1 had no effect on the growth of HCT-8 cells (**Fig. 1D**; bars 1a vs. 1b), but increased the tolerance to Roc-A (**Fig. 1D**; bars 2a vs. 2b). The treatment of elacridar, which is a selective MDR1 inhibitor, reduced the Roc-A tolerance of the host cells in a dose-dependent manner, but HCT-8/MDR1 cells were significantly more tolerant to Roc-A than HCT-8/NC cells (**Fig. 1D**; bars from 3a to 6b). These indicate that Roc-A is a native substrate of MDR1. Using this model, overexpressing MDR1 in host cells showed no effect on the anti-cryptosporidial efficacy of Roc-A (i.e., EC_50_ = 2.35 nM in HCT-8/MDR1 cells vs. EC_50_ = 2.30 nM in HCT-8/NC cells; p = 0.9586 by multiple comparison test) (**Fig. 1E**), which excludes the off-target effect of Roc-A.

After having confirmed the in vitro efficacy, we further tested the effects of Roc-A on the excysted sporozoites and early stage of development of *C. parvum*. In the “effect on sporozoite assay”, we determined that free sporozoites could survive for 2 h in PBS at 25 °C based on the qRT-PCR of parasite 18S rRNA (**Fig. 2A**). The 2-h window was used to evaluate the effect of Roc-A on the viability of excysted sporozoites, in which Roc-A displayed lower nanomolar inhibitory activity (EC_50_ = 54 nM) (**Fig. 2B**). The EC_50_ value is 30.5-fold higher than that obtained in 44-h infection assay described above (i.e., 54 nM vs. 1.77 nM). In comparison, the EC_50_ value of paromomycin (a commonly used positive control anti-cryptosporidial compound) on excysted sporozoites is 280.7 μM, which is >5,000-fold higher than that of Roc-A.

**Fig. 2.**
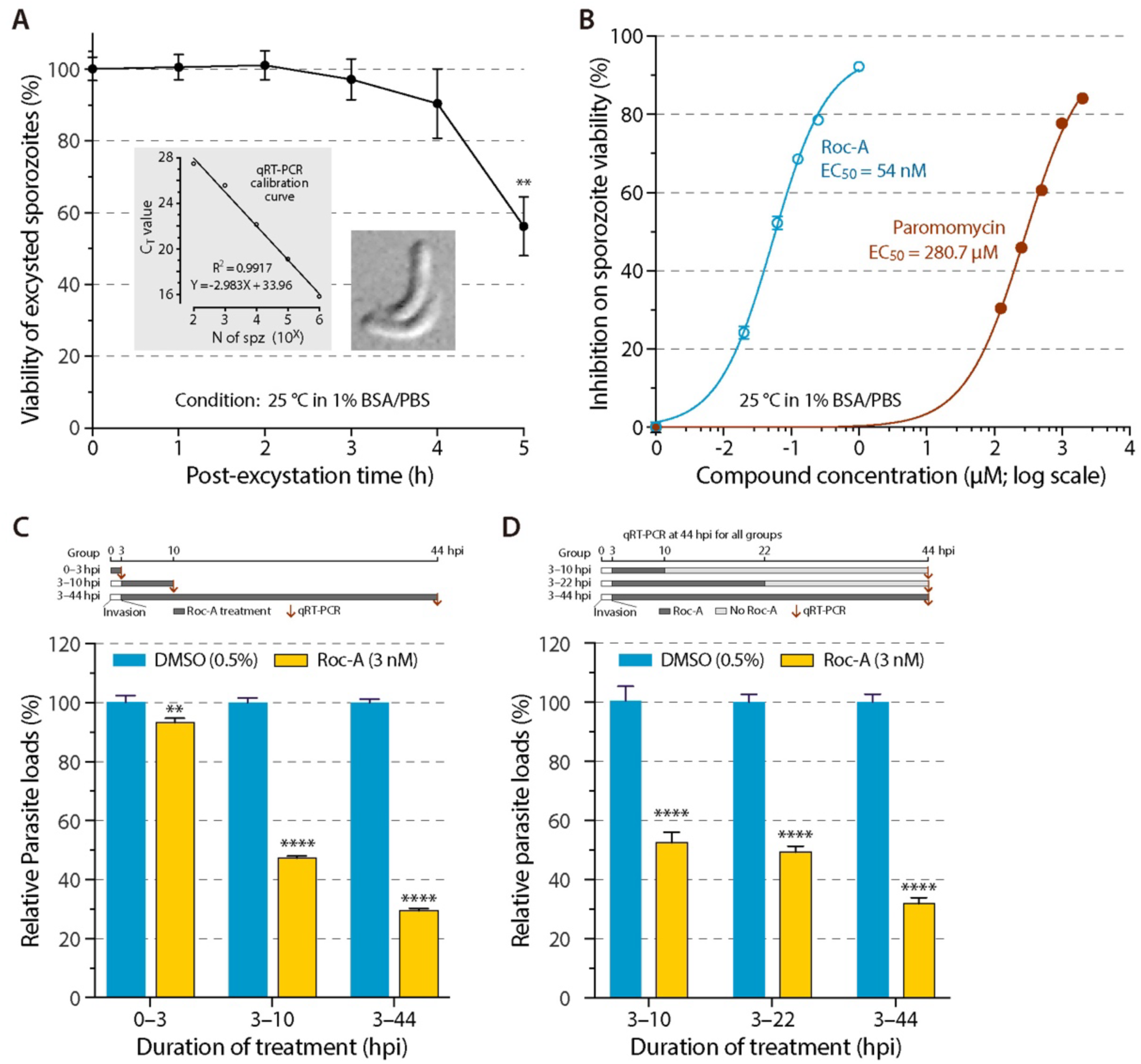
Effects of rocaglamide (Roc-A) on the viability of excysted sporozoites of *C. parvum* and on the parasite invasion and early stage development. **(A)** Viability of excysted free sporozoites at 25 °C in PBS (pH 7.4) containing 1% bovine serum albumin (BSA) over time. The viability was assessed by quantifying *C. parvum* 18S (Cp18S) rRNA by qRT-PCR. Insets show the qRT-PCR calibration curve with the C_T_ values against the number of sporozoites, and the morphology of two excysted sporozoites. **(B)** The viability of excysted sporozoites treated with Roc-A or paromomycin at specified concentrations in 1% BSA/PBS at 25 °C for 2 h by qRT- PCR detection of Cp18S rRNA. **(C)** Effect of Roc-A (3 nM in 0.5% DMSO; vs. 0.5% DMSO diluent control) on the parasite invasion (0–3 hpi treatment group) and first generation of merogony (3–10 hpi group) of *C. parvum* in comparison with a full-course treatment (3–44 hpi group). The parasite loads were assessed by the end of each treatment by qRT-PCR. Experimental design is illustrated on top of the panel. **(D)** Drug withdrawal assay to assess parasitistatic or parasiticidal effect of Roc-A. Invaded sporozoites were cultured in the presence of Roc-A for specified time periods, followed by the removal of Roc-A and continuous growth of *C. parvum* up to 44 h. Parasite loads were quantified by qRT-PCR at 44 hpi. Experimental design is illustrated on top of the panel. In (C) and (D), oocysts were added into the microplates containing HCT-8 monolayers, and allowed for excystation of sporozoites and invasion for 3 h (0–3 hpi), followed by the removal of uninvaded parasites and continuous culture for specified time periods. ** = p<0.01 and **** = p<0.0001 by Sidak’s multiple comparisons test (vs. 0.5% DMSO control at specified timepoints).

In the “effect on early stage assay” (**Fig. 2C**), *C. parvum* was treated with Roc-A at 3 nM during the invasion (0–3 hpi treatment group) and early stage development (3–10 hpi), together with a full course treatment (3–44 hpi). Parasite loads were quantified by qRT-PCR at the end of each treatment. In this assay, Roc-A (3 nM) slightly affected the parasite invasion (i.e., 6.7% reduction of parasite loads) (**Fig. 2C**, 0–3 hpi group), although the reduction is statistically significant (p<0.01 by Sidak’s multiple comparisons test). After the invasion, Roc-A (3 nM) treatment for 7 h already showed high efficacy, resulting in 52.6% reduction of the parasite loads (p<0.0001) (**Fig. 2C**, 3-10 hpi group). In the same assay, a full course treatment of Roc-A (3 nM) reduced the parasite load by 70.5% (P<0.0001) (**Fig. 2C**, 3–44 hpi group), which is comparable to the 62.7% reduction observed in the EC_50_ assay described above and depicted in **Fig. 1B**.

We also performed “drug withdrawal assay” to evaluate whether Roc-A was parasitistatic or parasiticidal for *C. parvum* by testing whether intracellularly developing *C. parvum* could recover from the early stage treatment with Roc-A (**Fig. 2D**). Post-invasion *C. parvum* was treated with Roc-A (3 nM) for 7 h (3–10 hpi) or 19 h (3–22 hpi), followed by the removal of Roc-A and continuous growth of *C. parvum* for up to 44 hpi, along with a full-course of treatment (3–44 hpi group). The parasite loads were quantified at 44 hpi in all groups. In this assay, the 7 and 19 h treatments reduced parasite loads by 47.4% and 50.5%, respectively (**Fig. 2D**). The full-course treatment (3–44 hpi) reduced the parasite loads by 68.1%, which is comparable to the 70.5% and 62.7% reductions in the two experiments described above (**Fig. 1B** and **Fig. 2C**). The 47.4% reduction in the 7-h treatment group in the “drug withdrawal assay” is close to the 52.6% reduction in the 7-h group in the “effect on early stage assay”, indicating that the growth of intracellularly developing *C. parvum* is unable to recover after the early inhibition by Roc-A. In other words, Roc-A is parasiticidal for *C. parvum*.

Collectively, we conclude that the selective eIF4A inhibitor Roc-A is highly efficacious and selective against the growth of *C. parvum* in vitro. The safety margin of Roc-A is remarkably high, and the anti-parasitic efficacy of Roc-A is attributed fully to its action on the parasite target. Roc-A displays a fast-killing parasiticidal against *C. parvum* in vitro.

### Roc-A is highly efficacious and acts fast in vivo against the chronic infection of *C. parvum* in IFN-γ KO mice

We then evaluated the in vivo anti-cryptosporidial activity of Roc-A in mice. The selection of Roc-A dose was first determined by a drug tolerance experiment, in which C57BL/6 mice were well tolerant to 7-day single daily oral dose of Roc-A at 0.5 mg/kg based on body weight gains and health scores (**S1 Fig**). In the drug efficacy experiment (see **Fig. 3A** for experimental design), chronic cryptosporidial infections were first established in IFN-γ KO mice (C57BL/6 background; N = 15; gender random) by a single inoculation of *C. parvum* oocysts (5×10^4^ oocysts/mouse). Peak oocyst shedding was detected by qPCR on 10 dpi (day post-infection) (**Fig. 3B** and **3C**). There was a decline of oocyst shedding after the peak in the subsequent days in the untreated control group (i.e. from 62.9% on 11 dpi to 51.1% on 15 dpi in the vehicle control group) (**Table 1**; **Fig. 3C**, blue lines). The fluctuation of oocyst productions over time is expected for both acute and chronic infection in animals.

**Fig. 3.**
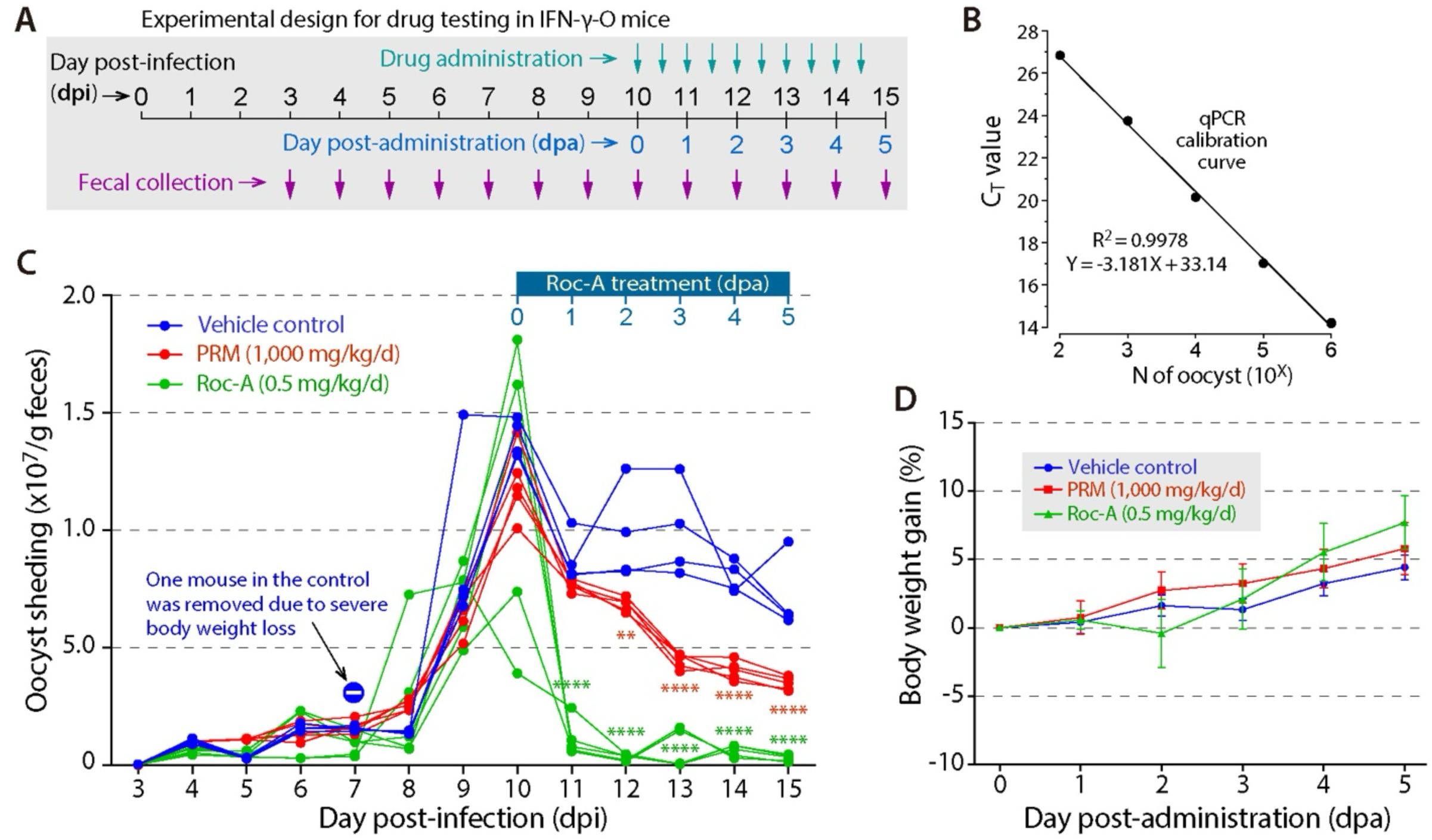
In vivo efficacy of Roc-A on the chronic infection of *C. parvum* in interferon-γ- knockout (IFN-γ-KO) mice. **(A)** Illustration of experimental design in the in vivo drug efficacy assay. **(B)** Standard qPCR curves derived by spiking varied numbers of *C. parvum* oocysts into *Cryptosporidium*-negative fecal samples, following the same DNA extraction and qPCR procedures as described for in vivo drug efficacy assay. **(C)** Effect of Roc-A (0.5 mg/kg/d in split doses) on the oocyst shedding in IFN-γ-KO mice chronically infected with *C. parvum* (N = 5/group). Untreated negative control used vehicle (1% DMSO). Treatment with paromomycin (1,000 mg/kg/d in split doses) was included for comparison. Fecal oocysts from individual mice were detected by qPCR and the quantities were obtained using the qPCR calibration curve shown in (B). ** = p<0.01 and **** = p<0.0001 by Tukey’s multiple comparisons test (vs. vehicle control on individual days). Note: one mouse in the control group was removed from later experiment due to severe body weight loss. **(D)** Percent body weight gains of mice chronically infected with *C. parvum* after the administration of Roc-A and paromomycin in comparison to the vehicle control. Bars indicate standard error of the mean (SEM). No statistical significances were detected on all individual days by Tukey’s multiple comparisons test (vs. vehicle control).

**Table 1.** Relative levels of fecal oocyst shedding in C57BL/7 interferon-γ knockout mice chronically infected with *Cryptosporidium parvum*. Abbreviations: DPA, day post-administration of compounds; DPI, day post-infection. DMSO, dimethyl sulfoxide; PRM, paromomycin; Roc-A, rocaglamide-A.

| Baseline for normalization | DPA | DPI | Vehicle (1% DMSO) |  |  | PRM (1000 mg/kg/d) |  |  | Roc-A (0.5 mg/kg/d) |  |  |
| --- | --- | --- | --- | --- | --- | --- | --- | --- | --- | --- | --- |
|  |  |  | Mean | SEM | N | Mean | SEM | N | Mean | SEM | N |
| In each group<br>(vs. 0 dpa) | 0 | 10 | 100.0 | 2.9 | 4 | 100.0 | 5.5 | 5 | 100.0 | 22.6 | 5 |
|  | 1 | 11 | 62.9 | 3.7 | 4 | 63.9 | 0.9 | 5 | 9.1 | 2.8 | 5 |
|  | 2 | 12 | 70.1 | 7.3 | 4 | 57.0 | 1.1 | 5 | 2.7 | 0.5 | 5 |
|  | 3 | 13 | 71.2 | 7.1 | 4 | 37.1 | 1.2 | 5 | 5.4 | 3.0 | 5 |
|  | 4 | 14 | 57.5 | 2.4 | 4 | 33.6 | 1.5 | 5 | 5.0 | 0.9 | 5 |
|  | 5 | 15 | 51.1 | 5.7 | 4 | 29.0 | 1.0 | 5 | 2.5 | 0.5 | 5 |
| On each day<br>(vs. vehicle) | 0 | 10 | 100.0 | 2.9 | 4 | 85.9 | 4.8 | 5 | 86.4 | 19.5 | 5 |
|  | 1 | 11 | 100.0 | 6.0 | 4 | 87.4 | 1.2 | 5 | 12.6 | 3.9 | 5 |
|  | 2 | 12 | 100.0 | 10.5 | 4 | 69.9 | 1.3 | 5 | 3.3 | 0.6 | 5 |
|  | 3 | 13 | 100.0 | 10.0 | 4 | 44.7 | 1.4 | 5 | 6.5 | 3.7 | 5 |
|  | 4 | 14 | 100.0 | 4.1 | 4 | 50.3 | 2.2 | 5 | 7.5 | 1.4 | 5 |
|  | 5 | 15 | 100.0 | 11.2 | 4 | 48.8 | 1.7 | 5 | 4.2 | 0.9 | 5 |

After oral administration of compounds on 0 day post-administration (dpa) (0 dpa = 10 dpi) (**Table 1**; **Fig. 3C**), Roc-A treatment (0.5 mg/kg/d; given in split doses at 0.25 mg/kg in 12 h intervals) resulted in a quick and remarkable decline of oocyst shedding (i.e., 90.9% to 97.5% reduction from 1 to 5 dpa; vs. 0 dpa). In comparison, the naturally occurring reductions of oocyst shedding in the vehicle group (1% DMSO) were between 37.1% (1 dpa) to 48.9% (5 dpa); while the treatment with paromomycin (PRM; 1000 mg/kg/d) produced moderate and gradual decline of oocyst shedding, from 36.1% (1 dpa) to 71.0% (5 dpa). In comparison with the vehicle group, the mouse body weight gains were slightly improved with PRM treatment, while those were slightly declined with the Roc-A treatment on 2 dpa but improved faster on the subsequent days (**Fig. 3D**). However, the differences between the three groups were not significant in any of the post-administration days by multiple comparison test. Overall, the in vivo data indicates that Roc-A is highly efficacious, and much more effective than PRM, against the cryptosporidial infection in mice. The >90% decline on the next post-administration day also indicates that Roc-A is a fast-killing anti-cryptosporidial agent, which is in congruent with the fast-killing action in vitro described above.

### Roc-A acts on the cytoplasmic translation initiation factor 4A in *C. parvum* (CpeIF4A) with lower nanomolar activity

The *C. parvum* genome encodes an eIF4A (gene ID cgd1_880), which contains domains and residues characteristic to eIF4A subfamily proteins, including an N-terminal DEAD-box helicase domain (InterPro name IPR044728) and a C-terminal helicase domain (IPR001650), as well as conserved residues for binding to ATP and interacting with eIF4G subunit (**Fig. 4A** and **4B**).

**Fig. 4.**
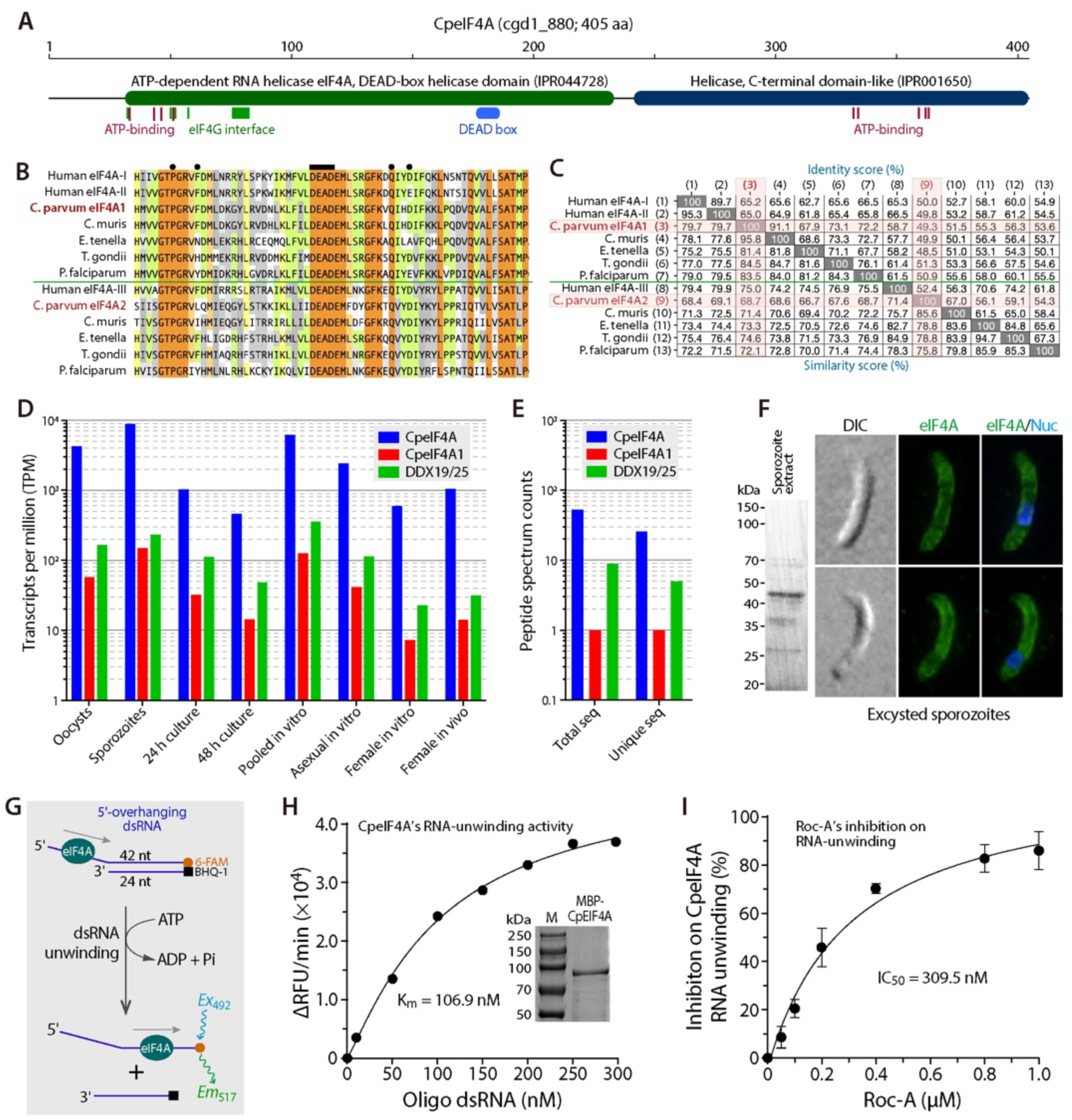
Molecular and biochemical features of *C. parvum* eukaryotic translational initiation factor 4A (CpeIF4A). **(A)** Domain architecture of CpeIF4A (cgd1_880; 405 aa) based on InterProScan analysis of domain families and functional residues for binding to ATP and interacting with eIF4G). **(B–C)** Comparison of amino acid sequence at the DEAD-box region (B) and identity/similarity scores (C) between CpeIF4A, human eIF4A-I/II (global initiation factors/RNA helicases) and orthologs from selected apicomplexan species (i.e., *Cryptosporidium muris, Eimeria tenella, Toxoplasma gondii* and *Plasmodium falciparum*), together with the closely related eIF4A-III-type orthologs (RNA helicase in exon junction complex), including CpeIF4A1, human eIF4A-III, and orthologs from selected apicomplexan species. In (B), the bar on top and dark circles indicate the DEAD-box (DDX) and the Roc-A binding sites based on the PDB model 5ZC9. **(D–E)** *CpeIF4A* gene expression levels in comparison with *CpeIF4A1* based on transcriptomic (D) and proteomic (E) data extracted from the CryptoDB, showing that *CpeIF4A1* is expressed at much higher levels in all developmental stages. Also see S1 Table and S2 Table for more detailed transcriptomic and proteomic data. **(F)** Immunostaining of CpeIF4A in sporozoites, showing cytoplasmic localization with stronger signals under the plasma membrane. Excysted sporozoites were immuno-stained with affinity-purified anti-CpeIF4A antibody (green) and counterstained with 4’,6-diamidino-2-phenylindole (DAPI) for nuclei (blue). **(G–H)** Biochemical features of CpeIF4A as determined by the RNA-unwinding assay (G), including enzyme kinetics towards the dsRNA (H) and inhibitory kinetics by Roc-A (I). Inset in (H) shows SDS-PAGE analysis of purified recombinant CpeIF4A.

*Cryptosporidium* also possesses a closely related gene annotated as eIF4A1 (cgd7_3940), together with 23 other DEAD-box (DDX) RNA helicases (total N = 25). These DDX RNA helicases contain fully conserved DEAD motif (N = 19) or partially conserved DExD or DEAx motifs (N = 5 or 1, respectively). Among them, CpeIF4A shows the highest identities to the human eIF4A-I/II, which are primarily involved in global translation initiation in the cytoplasm [28,33]; while CpeIF4A1 shows the highest identity to human eIF4A-III (**Fig. 4C**), which functions in the nucleus as a core component of the exon junction complex (EJC), playing a role in mRNA splicing [34,35]. To further determine the identities of CpeIF4A and CpeIF4A1, Bayesian inference (BI)-based phylogenetic analysis was performed for the DDX proteins from various *Cryptosporidium* and selected apicomplexan species with the highest sequence identities with human eIF4A-I/II and III, together with the orthologs from humans. The resulting BI tree separates these sequences into seven clusters (**S2 Fig**). CpeIF4A is clustered with human eIF4A-I and II, while CpeIF4A1 is grouped with human eIF4A-III.

Additionally, *CpeIF4A* gene is expressed at much higher levels than *CpeIF4A1* gene at various developmental stages in vitro based on the transcriptomic and proteomic datasets available at the CryptoDB, in which the expression of *CpeIF4A* (cgd1_880) is 31.9 to 82.1-fold higher in transcript abundance (transcriptomes), or 26 to 115-fold higher in peptide abundance (proteomes), than *CpeIF4A1* in various samples (**Fig. 4D** and **4E; S1** and **S2 Tables**). The cytoplasmic location of CpeIF4A is also confirmed by immunostaining using an anti-CpeIF4A polyclonal antibody produced in house (**Fig. 4F**), which shows that CpeIF4A is distributed in the cytoplasm of sporozoites, but the distribution is not homogenous and more signals are present under the plasma membrane. Similar cytoplasmic distribution pattern with stronger signals under the plasma membrane is also shown for intracellular developing *C. parvum* (**S3 Fig**). The cytoplasmic location, together with the high level of expression in all developmental stages and the phylogenetic affiliation with human eIF4A-I/II, supports that CpeIF4A is the housekeeping translation initiation factor in the parasite.

The RNA helicase activity of CpeIF4A is biochemically confirmed using a fluorescence-based RNA-unwinding assay (**Fig. 4G**). The recombinant CpeIF4A, which was expressed as a maltose binding protein (MBP)-fusion protein, displayed lower nanomolar activity to unwind dsRNA (K_m_ = 106.9 nM) (**Fig. 4H**). The RNA-unwinding activity of CpeIF4A could be specifically inhibited by Roc-A at sub-micromolar concentrations (IC_50_ = 309.5 nM) (**Fig. 4I**).

The binding affinity between Roc-A and CpeIF4A was confirmed using thermal shift assay (TSA), in which Roc-A displayed lower micromolar apparent dissociation constant (App. K_d_ = 3.51 μM) (**Fig. 5A**). The binding affinity was dramatically increased by over 100-fold in the presence of dsRNA and ATP (i.e., App. K_d_ = 33.7 nM) (**Fig. 5B**). For comparison, the binding affinity of dsRNA or ATP to CpeIF4A is at lower or higher micromolar levels (App. **K_d_** = 1.13 μM or 625.7 μM, respectively) **(Fig. 5C** and **5D**). These observations indicate that Roc-A binds much more tightly to the CpeIF4A-dsRNA-ATP complex than to CpeIF4A alone, which fits the structural model for human eIF4A-I [36]. In this model (PDB: 5ZC9), Roc-A clamps eIF4A onto polypurine sequences in mRNA via binding to the “bi-molecular cavity” formed by eIF4A-I and a sharply bent pair of consecutive purines in the RNA, thus blocking the translation initiation (**S4 Fig**). The observed, relatively lower affinity between Roc-A and CpeIF4A alone is in favor of the subsequent formation of much tighter binding of Roc-A to the CpeIF4A-RNA-ATP complex by making Roc-A more accessible to the bi-molecular cavity.

**Fig. 5.**
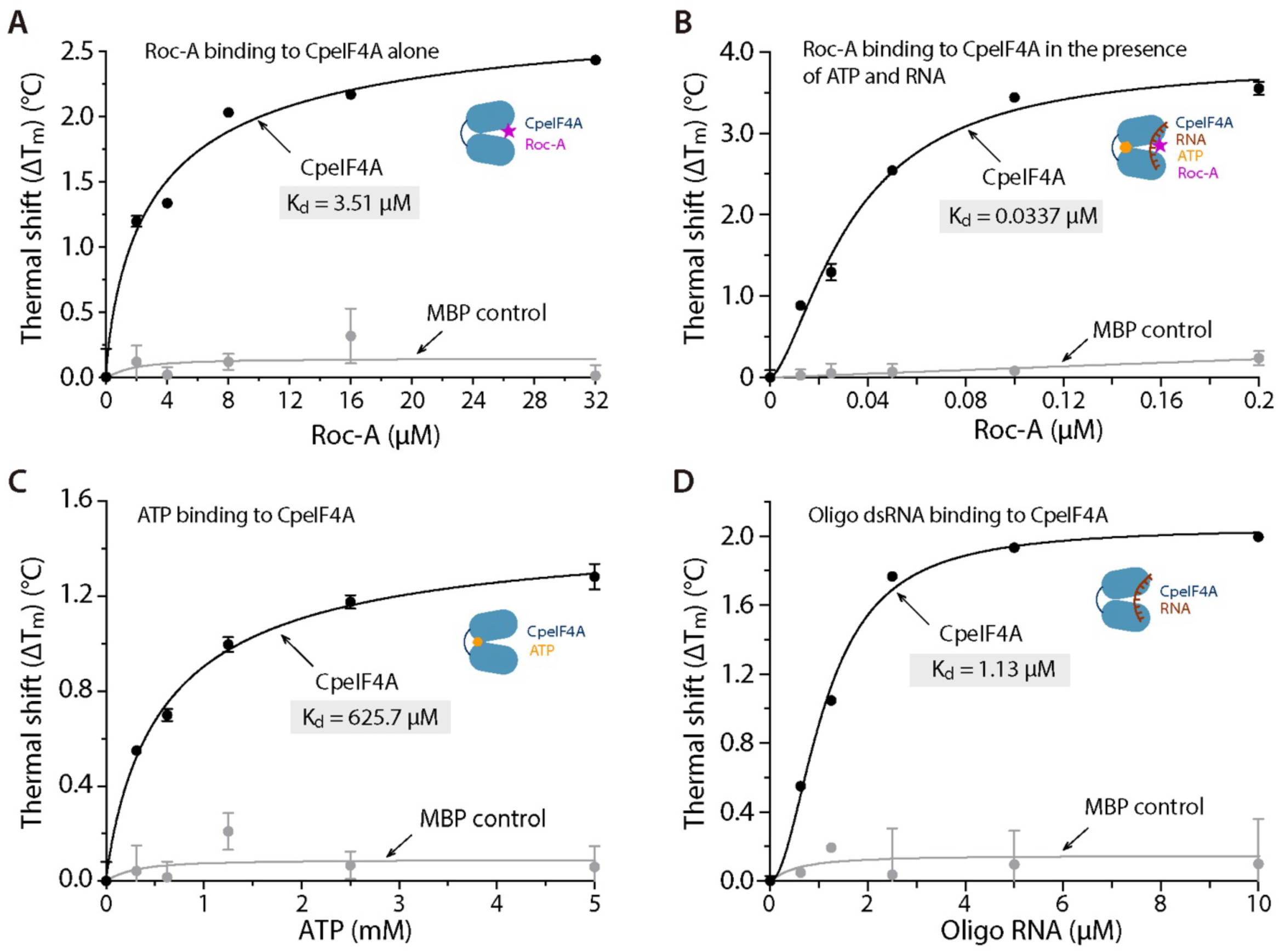
Binding affinities of CpeIF4A with Roc-A and substrates assessed by thermal shift assay (TSA). (A–B) Binding affinities of Roc-A with CpeIF4A alone (A) and in the presence of substrates dsRNA and ATP (B). The presence of dsRNA and ATP substantially increases the binding affinity (i.e., 104-fold lower K_d_ value). **(C–D)** Binding affinities of CpeIF4A with substrates, including ATP (C) and dsRNA (D).

In short, CpeIF4A is a cytoplasmic translational initiation factor with confirmed RNA helicase/RNA-unwinding activity. The helicase activity can be effectively inhibited by Roc-A via tight binding to the CpeIF4A-RNA-ATP complex.

### The inhibition of CpeIF4A by Roc-A results in the suppression of protein synthesis in excysting sporozoites of *C. parvum*, but affect little on the overall gene transcription

The inhibition of translation initiation by Roc-A would expectedly result in the repression of protein synthesis. This notion was tested by measuring the protein synthesis in *C. parvum* sporozoites that undergo excystation from oocysts, for which the pure parasite material could be practically obtained. In pilot experiments, we determined that Roc-A at concentrations up to 100 μM had no or little effect on the excystation and viability of excysted sporozoites (**Fig. 6A**), in which oocysts were incubated in excystation medium for 2 h, followed by the quantification of *C. parvum* 18S rRNA by qRT-PCR. While the excystation typically takes less than 1 h to complete, the 2 h incubation gives extra time for protein synthesis without affecting the sporozoite viability under host cell-free environment. Under this experimental condition, Roc-A treatment up to 10 μM also had no or little effect on the subcellular distribution of CpeIF4A in excysted sporozoites (**Fig. 6B**). The ineffectiveness of Roc-A on the parasite excystation is explainable, as the sporozoites in the intact oocysts are protected by the oocyst wall until the suture of oocysts opens; and once the suture opens, the exit of sporozoites is rapid (**Fig. 6F**). Nonetheless, this gives us a two-hour window to assess the effect of Roc-A on the protein synthesis in the excysting and excysted sporozoites.

**Fig. 6.**
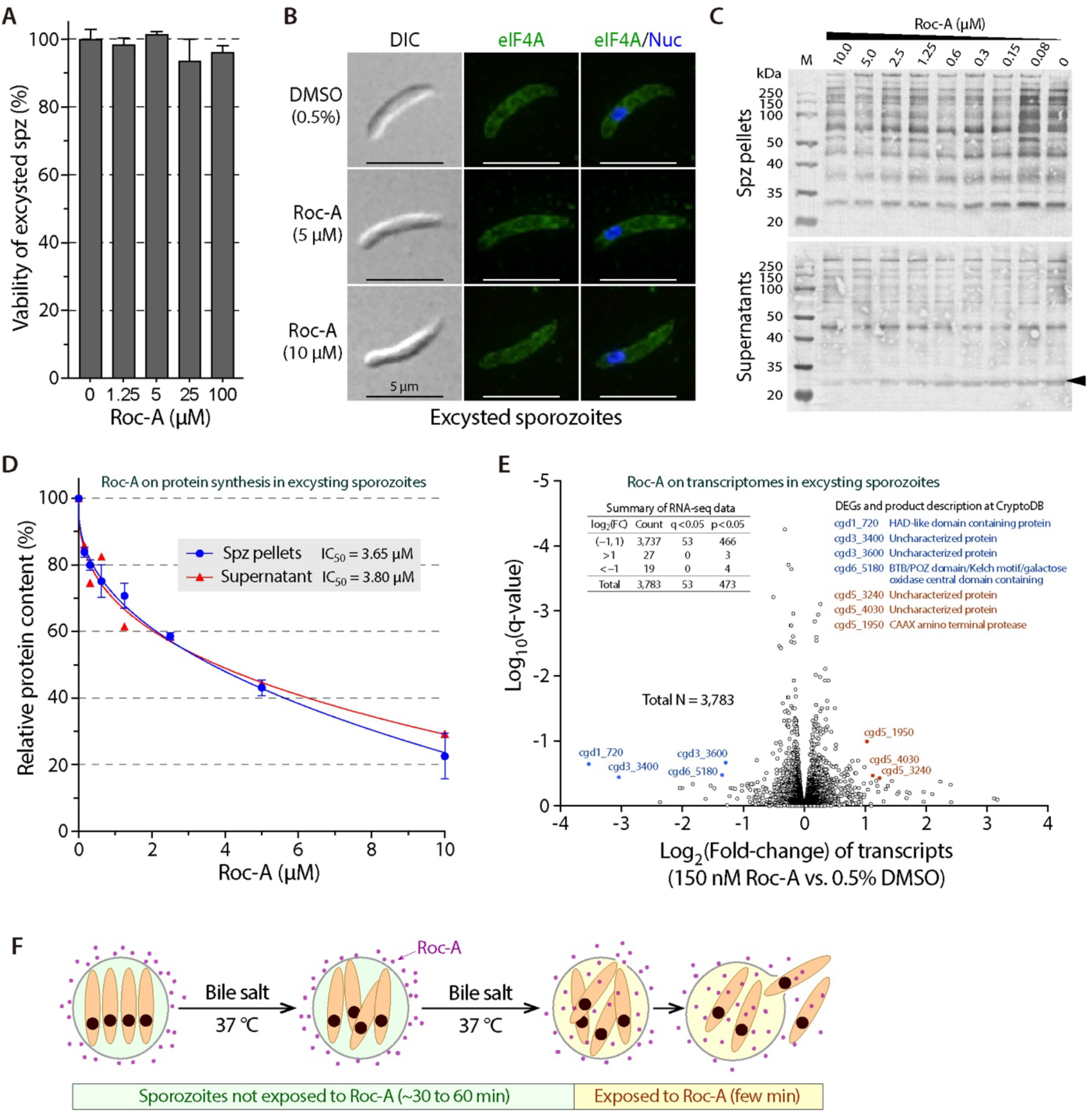
Effects of Roc-A on the translation and transcription in excysting sporozoites of *C. parvum*. **(A)** Effect of Roc-A on the viability of excysting *C. parvum* sporozoites, in which oocysts were allowed for in vitro excystation for 2 h in the absence or presence of Roc-A at varied concentrations. The data shows that *C. parvum* during the excystation process is tolerant to Roc-A up to 100 μM. **(B)** Effect of Roc-A at 5 and 10 μM on the morphology of sporozoites and subcellular distribution of CpeIF4A. Oocysts were treated as described above in (A), and excysted sporozoites were immuno-stained with affinity-purified anti-CpeIF4A antibody (green) and counterstained with 4’,6-diamidino-2-phenylindole (DAPI) for nuclei (blue). **(C)** Effect of Roc-A on the translation in sporozoites during excystation by western blot analysis. After excystation, sporozoite (Spz) pellets (representing non-secreted proteins) and supernatants (representing secreted proteins) were collected for blotting and immuno-stained with a pan-anti-*Cryptosporidium* polyclonal antibody. Most of the protein bands from the pellets (upper panel), but only one major band in the supernatant at 28 kDa (lower panel; marked by an arrowhead), showed decrease of intensities in response to Roc-A treatment. **(D)** Grayscale plot of the protein band intensities against the Roc-A concentrations based on the blots represented in (C). For non-secreted proteins (blue line), the intensities of total proteins (whole lanes) were measured; while for secreted proteins (red line), only the intensities of the 28-kDa protein were measured. Bars indicate standard error of the mean (SEM). **(E)** Effect of Roc-A on the transcriptomes in excysted sporozoites as determined by RNA-seq and shown by volcano plot. Among 3,783 detected gene transcripts, none of them are significantly changed using the criteria of |log_2_(Fold-change)| >2 and q-value <0.05. Only the transcripts of seven lowly expressed genes with undefined functions showed significant fold changes using a relaxed criteria of |log_2_(Fold-change)| >2 and p-value <0.05. **(F)** Illustration of the excystation process, showing that Roc-A has no access to the sporozoites in the first phase of excystation until the oocysts are ruptured and sporozoites start to escape from the oocysts.

We then evaluated the effect of Roc-A at concentrations up to 10 μM on the protein synthesis in excysting sporozoites by a quantitative western blot analysis using a rabbit antiserum raised against total protein extracted from *C. parvum* oocysts (pan-anti-*Cryptosporidium* antibody). The treatment of Roc-A resulted in dose-dependent reductions of protein band intensities in the excysted sporozoites (i.e., non-secreted proteins in sporozoite pellets) (**Fig. 6C**, upper), showing lower micromolar inhibitory activity based on grayscale plot of total proteins in individual lanes (IC_50_ = 3.65 μM) (**Fig. 6D**, blue line). On the other hand, there were no apparent reductions for the majority of the secreted proteins in the supernatants (**Fig. 6C**; lower), but only one of the major bands at 28 kDa showed significant reduction (**Fig. 6C**; arrowhead). This 28-kDa band also displayed a dose-dependent curve and IC_50_ value that are similar to those for the non-secreted proteins (IC_50_ = 3.80 μM) (**Fig. 6D**, red line).

The similarity of inhibitory kinetics between the non-secreted proteins and the single 28-kDa secreted protein indicates that the synthesis of new proteins in sporozoites during the excystation are CpeIF4A-dependent and subjected to the inhibition by Roc-A. While the antiserum may not recognize all proteins present in excysted sporozoites, our data indicates that much more non-secreted proteins are newly synthesized than secreted proteins during the excystation process. It agrees with the notion that most of the secreted protein would have been already synthesized and stored in sporozoites before sporozoites start to escape from the ruptured oocysts. Nonetheless, our data support the model that the translation initiation machinery can be blocked by Roc-A via acting on CpeIF4A, thus suppressing the protein synthesis in the parasite.

In contrast to the suppression on protein synthesis, Roc-A treatment produces little effect on the transcriptome in sporozoites (**Fig. 6E**). In this experiment, free sporozoites after excystation were treated with Roc-A or 0.5% DMSO diluent (three biological replicates) at 150 nM for 2 h at 25 °C, which corresponds to the EC_70_ concentration as described above and depicted in **Fig. 2B**. RNA-seq detected the transcripts of 3,795 protein-coding genes, in which the transcripts of 3,783 genes had one or more reads in either Roc-A-treated or the 0.5% DMSO control groups.

However, no differentially expressed genes (DEGs) were observed using q-value (aka. adjusted P-value) of 0.05 and 2-fold change (FC) as the cutoffs (**Fig. 6E**). Using a more relaxed statistical significance cutoff (i.e., unadjusted p = 0.05 and |log2FC| >2 as cutoffs), only seven DEGs (0.185%) were identified, including four down-regulated and three up-regulated (**Fig. 6E**). All seven genes are of unknown functions and lowly expressed (relative abundances from 7.5 to 32.4 in the control and from 0.7 to 56.2 in the Roc-A-treated groups; vs. median values of 617.2 and 625.0, and mean values of 4,612.3 and 4,485.3 in the control and treated groups, respectively) (**Fig. 6E; S3 Table**), including four encoding uncharacterized proteins and three containing certain domains for which biological roles cannot be assigned.

## Discussion

This study provides a chain of evidence that the translational machinery can be targeted via inhibition of the eukaryotic elongation factor 4A (eIF4A) for developing fast-killing anti-cryptosporidial therapeutics. We show that Roc-A, a selective and specific inhibitor of eIF4A, is highly efficacious against the growth of *C. parvum* in vitro (EC_50_ = 1.77 nM) and in vivo (>90% reduction of oocyst shedding at 0.5 mg/kg/d in mice on the second day post-administration). Using an in vitro model over-expressing MDR1, we confirm that the killing of the intracellularly developing *C. parvum* by Roc-A is fully attributed to the action on the parasite target (on-target effect). Roc-A acts fast on *C. parvum*, and the killing is irreversible (parasiticidal). Roc-A inhibits the helicase/RNA-unwinding activity of CpeIF4A (IC_50_ = 309.5 nM), the global eukaryotic transcriptional factor 4A in the parasite cytoplasm, via tight binding to the CpeIF4A-RNA-ATP complex (K_d_ = 33.7 nM). The action of Roc-A on CpeIF4A suppresses in the protein synthesis in the excysting sporozoites, mainly on the non-secreted proteins (IC_50_ = 3.65 to 3.80).

The single-digit lower nanomolar in vitro efficacy and extremely high selectivity (SI >1,400) suggests that *C. parvum* is much more susceptible to the inhibition of protein synthesis than host cells. It is likely that Roc-A also acts on other DDX proteins in *C. parvum*, such as CpeIF4A1, resulting in a synergistic killing effect on the parasite, as Roc-A has also known to target human DDX3 albeit with lower affinity [37]. While the in vitro selectivity of Roc-A is extremely high (SI >1,400), the safety margin on mice is relatively low (certain toxicity observed at 1.5 mg/kg/d; vs. well-tolerated at 0.5 mg/kg/d), suggesting that certain types of cells in animals are more sensitive to Roc-A than enterocytes and hepatocytes. Nonetheless, our data validate that CpeIF4A and its orthologs in other *Cryptosporidium* species can serve as a novel drug target for developing inhibitors with better in vivo selectivity. Roc-A as an anti-cryptosporidial lead is also worth being explored for the potential as anti-cryptosporidial drug, as well as for lead optimization. Additionally, the high efficacy of Roc-A on the sporozoites and intracellular *C. parvum* in vitro indicates that this inhibitor can serve as a valuable probe for the investigation of the yet poorly understood mechanism and regulation of protein synthesis in the parasite.

It is less expected, but intriguing, that the inhibition of CpeIF4A by Roc-A can lead to the suppressive effect on the protein synthesis in excysting sporozoites, but not on the transcription (or the effect on the transcriptomes is insignificant). It is likely attributed to the “delayed effect” from the suppression of protein synthesis to the inhibition of transcription. In other words, excysted sporozoites may use existing translational machinery consisting of various proteins that have been synthesized prior to the exposure to Roc-A, by which significant effect of Roc-A on the transcription can be observed only after the existing transcriptional factors are depleted. This is more obvious during the excystation process (**Fig. 6F**), in which the sporozoites stay in intact oocysts without exposure to Roc-A for a relatively long time (usually 30–60 min), until the oocyst wall ruptures at the suture and sporozoites start to escape from the oocysts. The release of sporozoites from ruptured oocysts is quick, usually taking a few minutes. In addition to the delayed effect, it is also plausible that sporozoites after excystation may synthesize significant new proteins but only limited amount of new RNA species. While these notions need to be experimentally validated, they appear to be biologically justifiable, as sporozoites after excystation are present in free form for a very short period before they reach and invade host enterocytes (i.e., in minutes) [38], for which there is a little demand for them to synthesize significant amounts of new mRNA and/or proteins.

## Materials and methods

### Parasite and host cells

An isolate of *C. parvum* (gp60 subtype IIaA17G2R1) was propagated in calves and purified from calf feces by sucrose and CsCl gradient centrifugation protocols as described [39,40]. Purified oocysts were stored in PBS containing 200 U/mL penicillin and 0.2 mg/mL streptomycin at 4 °C until use. The viability of oocysts was examined regularly. Only those with >80% excystation rates were used in experiments. Before infection of host cells, oocysts were treated with 10% bleach in water for 5 min on ice, washed five times with PBS and resuspended in culture medium. HCT-8 or HepG2 cells were cultured with RPMI-1640 medium or DMEM medium, respectively, containing 10% fetal bovine serum (FBS), 50 units/mL penicillin, and 50 µg/mL streptomycin at 37 °C under 5% CO_2_ atmosphere.

### In vitro drug efficacy assay on *C. parvum*

The efficacy of Roc-A on the growth of *C. parvum* in vitro used a previously developed “44-h infection assay” with slight modifications, which included 3 h excystation/invasion and 41 h intracellular development in the presence of the compound at varied concentrations [29,30].

Briefly, HCT-8 cells were seeded in 96-well cell culture microplates and allowed to grow overnight until ∼80% confluence. The oocysts of *C. parvum*, which were bleach-treated and extensively washed, were suspended in RPMI-1640 medium containing 0.15% taurocholic acid and added into the microplate by a medium exchange (5×10^4^ oocysts/well). Inoculated oocysts were allowed for excystation and invasion for 3 h. Uninvaded parasites were removed by an exchange of RPMI-1640 medium containing 10% FBS, 50 units/mL penicillin, and 50 µg/mL streptomycin. At this timepoint, Roc-A at specified concentrations were added into the microplates, including diluent control (0.5% dimethyl sulfoxide [DMSO]). Parasite-infected monolayers were incubated at 37℃ in a 5% CO_2_ incubator for 41 h, followed by preparation of cell lysates and detection of parasite loads by qRT-PCR as described below.

At 44 h post-infection (hpi) timepoint, microplates were centrifuged and supernatants were removed. Ice-chilled bovine serum albumin (BSA; 0.1% solution prepared in pyrogen/nuclease-free water) was then added into the microplates (50 μL/well), followed by the vigorous vortexing on ice for 30 min as described [29,30]. After centrifugation (2,000 g for 15 min), supernatants (cell lysates) were used immediately for qRT-PCR or stored at −80 °C. The parasite loads were evaluated by qRT-PCR using HiScript II One Step qRT-PCR SYBR Green Kit (Vazyme, Nanjing, China). Reactions were performed in StepOnePlus Real-Time PCR System (Applied Biosystems, Carlsbad, CA). Each reaction in 20 μL final volume contained 2 μL cell lysate (100× dilution), 0.4 μL forward and reverse primers (each at 10 μM final concentration), 1 μL One Step SYBR Green Enzyme Mix, 0.4 μL 50× ROX reference dye I, 10 μL 2× One Step SYBR Green Mix, and 5.8 μL nuclease-free water.

The following primers were used to quantify the 18S rRNA levels for *C. parvum* (Cp18S) and host cell (Hs18S) as described [29,30]: Cp18S-1011F (5’-TTG TTC CTT ACT CCT TCA GCA C-3’) and Cp18S-1185R (5’-TCC TTC CTA TGT CTG GAC CTG-3’) and Hs18S (Hs18S-1F, 5’- GGC GCC CCC TCG ATG CTC TTA-3’ and Hs18S-1R, 5’-CCC CCG GCC GTC CCT CTT A-3’). The inhibition of parasite growth was calculated based on ΔΔC_T_ values, in which EC_50_ values were calculated using a sigmoidal model fit as described [29,30]. Standard curves were obtained by inoculating HCT-8 cells with varied numbers of oocysts, followed by the same 44-h infection and qRT-PCR procedures as described above.

### In vitro cytotoxicity assay

The cytotoxicity of Roc-A was evaluated by an MTS cell viability assay (Saint-Bio, Shanghai, China). HCT-8 and HepG2 cells (10^4^ cells/well) were seeded into 96-well microplates and culture overnight (16 h). Serially diluted Roc-A, including negative control with diluent (0.5% DMSO), was added into microplates and continuously cultured for 41 h. The plates were rinsed with PBS for three times, and incubated with 10 μL/well of MTS solution (37 °C for 2 h). The optical density at 490 nm (OD_495_) were measured in Synergy LX multi-mode reader (BioTek, Winooski, VT, USA). The 50% toxicity concentrations (TC_50_) on HCT-8 or HepG2 cells were calculated based on the OD_495_ values (TC_50_ = [OD_495(treat)_ – OD_495(blank)_]/[OD_495(ctl)_ – OD_495(blank)_]), in which OD_495(treat)_ and OD_495(ctl)_ refer to the groups treated with Roc-A at a specified concentration and 0.5% DMSO diluent, respectively; while OD_495(blank)_ refers to the read from the blank wells. Each group included three or more biological replicates. Based on the TC_50_ and EC_50_ values, in vitro selectivity index (SI) was calculated (SI = TC_50_/EC_50_).

### Drug effect on the excysted sporozoites and early intracellular developmental stages in vitro, and drug withdrawal assay

Effects of Roc-A on *C. parvum* sporozoites, invasion and early developmental stages were also evaluated using qRT-PCR assay. Free sporozoites were prepared by incubating oocysts with PBS (pH 7.4) containing 0.75% taurocholic acid at 37 °C for 45 min (>90% excystation rate), followed by centrifugation and resuspension of excysted sporozoites in PBS or RPMI-1640 medium. Since excysted sporozoites may survive only for a limited time under host cell-free condition, we evaluated the viability of free sporozoites by qRT-PCR, in which excysted sporozoites were resuspended in PBS containing 1.0 mg/mL BSA and divided into 18 aliquots (8×10^5^ sporozoites each). After incubation at 25 °C for 0, 1, 2, 3, 4 or 5 h (3 aliquots/group), samples were centrifuged and pellets were lysed in 50 μL of 0.1% BSA/nuclease-free water as described above. The viability of sporozoites over time was assessed by qRT-PCR detection of Cp18S rRNA. After having determined that free sporozoites could maintain viability at lease for 2 h (**Fig. 2A**), we used the 2 h window to test the effect of compounds on the viability of free sporozoites. Briefly, excysted sporozoites in 1% BSA/PBS were treated with serially diluted Roc-A or paromomycin containing 0.5% DMSO (including diluent control) at 25 °C for 2 h. Free sporozoites were collected by centrifugation and lysed in 0.1% BSA/nuclease-free water for qRT-PCR detection of Cp18S and calculation of EC_50_ values as described above. Standard curves were obtained using serially diluted sporozoites, followed by the preparation of lysates and quantitation of Cp18S rRNA by qRT-PCR.

Effect of Roc-A on the early intracellular development of *C. parvum* was tested similarly as for the 44-h infection assay, in which HCT-8 cell monolayers at ∼80% confluence were treated with Roc-A at 3 nM, which was close to the EC_65_ concentration obtained in the 44-h infection assay as depicted in Fig. 1B, during the excystation/invasion stage (0–3 hpi treatment group) or first generation merogony (3–10 hpi group). A full course 44-h infection assay (3–44 hpi group) was included for comparison. Cells were lysed at the end of each treatment. The parasite loads were quantified by qRT-PCR (see illustration in Fig. 2C).

Drug withdrawal assay was used to determine whether Roc-A is parasitistatic or parasiticidal for *C. parvum*, in which intracellular parasites after invasion were incubated with Roc-A (3 nM) between 3–10 hpi and 3–22 hpi. At the end of each treatment, Roc-A was removed by a medium exchange (drug withdrawal). The parasites were continuously cultured for up to 44 hpi. A full-course treatment group (3–44 hpi) was included for comparison (see illustration in Fig. 2D). The parasite loads for each group were determined at 44 hpi by qRT-PCR. In all in vitro assays, each group included three or more biological replicates.

### Validation of on-the-parasite-target effect of Roc-A using host cells transiently over-expressing human *MDR1* gene

The on-target effect of Roc-A was assessed using a recently developed in vitro model, which tested whether the increase of drug tolerance in host cells by over-expressing the human *MDR1* gene affected the drug efficacy on the epicellular *C. parvum* [31,32]. In this study, we first confirmed that Roc-A was a native substrate of MDR1, in which transient over-expression of *MDR1* in HCT-8 cells increased the drug tolerance to Roc-A in host cells (i.e., making host cells more susceptible to the inhibition of Roc-A in the presence of elacridar, a potent third generation of MDR1 inhibitor) (Fig. 1D). The transient over-expression of MDR1 in host cells was established by transfecting HCT-8 cells with pCVM3 that carried *MDR1* gene (pCMV3-MDR1), together with a blank vector control as described [32]. The anti-cryptosporidial efficacy of Roc-A in HCT-8 cells, which over-expressed MDR1 (HCT-8/MDR1) or blank vector negative control (HCT-8/NC), were performed as described above for the 44-h infection assay. The “on-the-parasite target” effect was assessed by comparing the inhibitory curves and EC_50_ values against the growth of *C. parvum* cultured with HCT-8/MDR1, HCT-8/NC and wild-type HCT-8 cells. No or little effect indicates that the anti-cryptosporidial activity was attributed fully to the drug action on the parasite target [31,32].

### In vivo drug efficacy assay on *C. parvum*

In vivo drug efficacy was evaluated in a mouse model of chronic infection of *C. parvum* using interferon-γ knockout (IFN-γ-KO) mice with C57BL/6 genetic background, which were housed in a double-layer isolator control system (Hebei Linhai Metal Products, Baoding, China). We first evaluated the tolerance of IFN-γ-KO mice on Roc-A, in which 12 mice (8-wk old; gender random) were randomly assigned into four groups (3 mice/group) for administration of Roc-A at 0, 0.5, 1.0 and 2.0 mg/kg/d for seven days (split into two doses given in 12 h internal). Roc-A was prepared in 0.5% DMSO in distilled water (vehicle), and the negative control was given with vehicle. Mice were weighed daily from 0 to 7 day post-administration (dpa). Mice with >20% weight losses (if any) were terminated from further observations by euthanasia. Mice were also observed and scored for fur condition, weight loss, hunchbackedness, and behavior. A health score was calculated for each mouse based on a previously reported scales with modifications as detailed (**S3 Table**) [41,42].

Based on the drug tolerance data, a split daily dose at 0.25 mg/kg (= 0.5 mg/kg/d) was used to evaluate the anti-cryptosporidial efficacy. A total of 15 IFN-γ-KO mice (8-week old; gender random) were orally inoculated with *C. parvum* oocysts (5×10^4^ oocyst/mouse) and allowed to establish chronic infection for 10 days. On 10 dpi (day post-infection), mice were randomly assigned into three groups for administration by oral gavage of Roc-A (0.5 mg/kg/d), paromomycin (1,000 mg/kg/d), or vehicle (0.5% DMSO in water), all at split doses in 12 h intervals, for 5 days. Mice were weighed daily from 0 to 5 dpa (day post-administration of compounds), corresponding to 10 to 15 dpi, in each morning prior to the first administration, plus a final collection at 24 h after the last administration of compounds. Fecal pellets were collected daily from 3 to 15 dpi in the mornings, plus a final collection at 24 h after the last administration of compounds, for which individual mice were placed in divided cardboard boxes for ∼1 h until sufficient fecal pellets were collected. Mice were also observed for health signs as described above in the drug tolerance experiment. The experiment was terminated by euthanasia of mice on 17 dpi. The experimental design is illustrated in Fig. 3A.

### Quantification of *C. parvum* oocyst shedding in mice

Oocyst shedding was quantified by qPCR. Fecal pellets from individual mice were weighed and soaked in equal volume of PBS for at least overnight at 4 °C. Fecal DNA was isolated using TIANamp Stool DNA Kit (Tiangen biotech, Beijing, China), in which 200 μL of homogenized fecal suspensions were mixed with equal volume of lysis buffer and subjected to six freeze-and-thaw cycles to break oocyst walls. The remaining procedures followed the manufacturer’s instruction. At the final step, DNA was eluted into 50 μL elution buffer. Fecal oocysts were qualified by qPCR detection of Cp18S rDNA using the same pair of primer as described above in the qRT-PCR-based in vitro efficacy assay. Standard curves were obtained by spiking fecal samples from *Cryptosporidium*-negative mice with serially diluted oocysts (100 to 10^6^ oocysts in 200 μL sample), followed by the sample procedures for the isolation of DNA and qPCR. The quantities of oocysts in experimental mice were calculated by calibration to the standard curves.

### Molecular analysis of CpeIF4A and phylogenetic reconstruction

The *C. parvum* genome encodes an eIF4A ortholog annotated at the CryptoDB as “Eukaryotic initiation factor 4A” (cgd1_880) and a closely related ortholog “eIF4A-1 eukaryotic translation initiation factor 4A-1 RNA SFII helicase” (cgd7_3940). In comparisons to human sequences, cgd1_880 and cgd7_3940 products share the highest identities to eIF4A-I/II and eIF4A-III, respectively. InterProScan analysis was performed on both gene products to identify domains and residues (https://www.ebi.ac.uk/interpro/search/sequence/). BLASTP searches were performed to identify closely related sequences from the NCBI’s protein databases, in which cgd1_880 and cgd7_3940 products showed highest identities to eIF4A-I/II (global transcriptional factor) and eIF4A-III (specialized function such as in mRNA splicing) subfamily proteins, respectively.

Bayesian inference (BI)-based phylogenetic analysis was performed for eIF4A-I/II and III orthologs, together with other closely related DDX proteins, from various *Cryptosporidium* species, selected apicomplexan species, and humans. Note: this phylogenetic analysis is not for elucidating the evolutionary history of *Cryptosporidium* eIF4A-related DDX proteins, but for assigning them into appropriate DDX protein subfamilies, for which the taxonomic sampling is limited. These sequences were fetched from NCBI’s reference proteins by BLASTP searches.

Multiple sequence alignments were performed using MUSCL (v3.8.31) (http://www.drive5.com/muscle/). After the removal of identical sequences and incomplete sequences, as well as gaps and ambiguously aligned positions, a final dataset consisting of 77 taxa and 335 positions were used for phylogenetic reconstructions using MrBayes (v3.2.7).

Amino acid substitution used mixed model, with the consideration of invariable sites and a four-rate gamma distribution for the variable sites. A total of 1,000,000 generations of searches with four independent chain running were performed, in which trees were sampled at an interval of every 1000 searches. Consensus tree was built by a majority ruling rule from sampled trees after the exclusion of first 25% trees.

### Expression of recombinant CpeIF4A and biochemical assays

A fragment of DNA encoding CpeIF4A (cgd1_880) was synthesized with codon optimized for expression in *Escherichia coli* (Sangon Biotech, Shanghai, China) and cloned into the pMAL-c5x vector for the expression of maltose-binding protein (MBP)-fused recombinant protein (MBP-CpeIF4A) in the BL21(DE3) strain of *E. coli*. The expression was induced by isopropyl-beta-D-thiogalactopyranoside (IPTG; 0.4 mM) at 16 °C for ∼16 h. MBP-CpeIF4A was affinity-purified using amylose resin and eluted with maltose solution as instructed by the manufacturer (New England Biolabs, MA, USA). Purified MBP-CpeIF4A was assessed by SDS-PAGE (10% gel), and quantified by Bradford protein assay using BSA as the standard. MBP-tag was similarly expressed from pMAL-c5x blank vector and purified for use as a negative control in the biochemical assays described below.

The helicase activity of MBP-CpeIF4A was detected by fluorescence resonance energy transfer (FRET)-based dsRNA-unwinding assay as illustrated in **Fig. 3G**. The 5’-overhanging dsRNA was synthesized by Sangon Biotech (Shanghai, China), which contained a long oligoribonucleotide (5’-CAU UAU CGG AUA GUG GAA CCU AGC UUC GAC UAU CGG AUA AUC-3’-6-FAM; 42 bases) complemented with a short one (BHQ-1-5’-GAU UAU CCG AUA GUC GAA GCU AGG-3’; 24 bases) [43]. The 5’-end overhang is for initial binding of eIF4A, and the BHQ1 (Black Hole Quencher 1) at the 3’-end quenches the emission from the 6- FAM (fluorophore 6-carboxyfluorescein) at the 5’-end of the other strand. When the dsRNA is unwound by CpeIF4A, the BHQ-1 separates from 6-FAM, by which the emission from 6-FAM became detectable (**Fig. 3G**). The assay (50 μL final volume) contained ATP (5 mM or as specified), RNasin ribonuclease inhibitor (20 U) and dsRNA (50 pmol or as specified), which were premixed into a 50 μL reaction buffer (50 mM HEPES, 2 mM DTT, 2 mM MgCl_2_ and 100 mM NaCl; pH 7.5) and were prewarmed to 37 °C. The reactions were started by adding 30 pmol MBP-CpeIF4A at the final step. MBP-tag (30 pmol) was used as negative control and for background subtraction. The RNA unwinding activity was detected in StepOnePlus Real-Time PCR System (Applied Biosystems, Carlsbad, CA; excitation at 492 nm and emission at 517 nm) at 37 °C and read by minute for 100 min. Further evaluation for the inhibitory effects of Roc-A on the dsRNA helicase activities as described above. Inhibitory kinetic curves were obtained by adding Roc-A at varied concentrations, from which 50% inhibition concentrations (IC_50_ values) were determined by four-parameter sigmoidal model [Y = (IC_max_ × X^h^)/(IC_50_^h^ + X^h^)].

### Binding affinity of CpeIF4A to substrates and Roc-A by thermal-shift assay (TSA)

The binding affinity was evaluated by thermal-shift assay (TSA) [44,45]. TSA assay was performed in microplates in 50 μL HEPES buffer (90 mM Na_2_HPO_4_, 10 mM NaH_2_PO_4_, 2 mM DTT; pH 7.8) containing MBP-eIF4A (5 μM) and SYPRO Orange (20 μM) in the absence or presence of ATP, dsRNA and/or Roc-A at specified concentrations. Microplates were briefly vortexed, centrifuged (10,000 g for 2 min), incubated at room temperature for 2 min, and placed in the real-time PCR system. Melting curves were obtained by gradual increase of temperature from 40 to 70 °C (1 °C/min). Based on the melting curves, melting temperatures (T_m_) were calculated using Boltzmann curve fit (Y = Bottom + (Top – Bottom)/(1 + EXP((T_m_ – X)/Slope))). Dissociation constants (K_d_ values) were calculated by Sigmoidal curve fit between the thermal shift (ΔT_m_) and the concentrations of substrates or Roc-A.

### Detection of native CpeIF4A by western blot analysis and immunofluorescence assay (IFA)

#### Antibody production

A rabbit anti-CpeIF4A pAb was produced against a synthetic peptide (^20^GEIESNYDEIVEC^32^) unique to CpeIF4A. A specific pathogen-free rabbit were immunized five times with keyhole limpet hemocyanin (KLH)-linked peptide, including 100 μg with Freund’s complete adjuvant for the first intradermal injection, 50 μg with incomplete adjuvant for next three intradermal injections, and 20 μg without adjuvant in the final intravenous injection.

Antiserum was collected and subjected to an affinity purification using the antigen immobilized to nitrocellulose membrane as described [46,47]. Briefly, 100 μg peptide antigen was dissolved in 300 μL pyrogen-free water, incubated with a nitrocellulose membrane (∼1 cm^2^) for 1 h, dried at 37 °C for 2 h, and blocked with 5% defat milk in TBST buffer (50 mM Tris-HCI at pH 7.5, 150 mM NaCl, and 0.05% Tween-20 for 1 h. After five washes with TBST buffer, the membrane was incubated with 10% antiserum or pre-immune serum in TBST (2 mL) for overnight at 4 °C, washed with TBST for five times, and eluted with 0.5 mL glycine at 200 mM prepared with 150 mM NaCl and 0.05% Tween-20 (pH 3.47). Eluted antibody was immediately neutralized with 8 μL 1M Tris-base to final pH 7.0.

#### Western blot analysis

Free sporozoites were prepared by in vitro excystation of *C. parvum* oocysts as described above and were lysed in RIPA buffer containing a protease inhibitor cocktail (Sigma-Aldrich). Sporozoite lysates (∼7.0×10^7^/lane) in reducing sample buffer were heated at 95 °C for 5 min, followed by SDS-PAGE and blotting onto nitrocellulose membranes. The blots were blocked in 5% BSA/TBST for 1 h and probed in 5% BSA/TBST containing affinity purified anti-CpeIF4A antibody (1:5 dilution) for 1 h. After three washes with TBST, blots were incubated with horseradish peroxidase-conjugated goat anti-mouse IgG (Invitrogen, Waltham, MA, USA) and visualized using an enhanced chemiluminescence reagent (Beyotime Biotechnology, Shanghai, China). All procedures were conducted at room temperature unless specified.

#### Immunofluorescence assay (IFA)

Free sporozoites were prepared by in vitro excystation with or without Roc-A at specified concentrations, fixed with cold paraformaldehyde (4%) for 1 h, washed three times in PBS, placed onto poly-L-lysine-coated slides and allowed to settle on the slides for 1 h. Sporozoites on the slides were permeabilized with Triton-X-100 (0.2% in PBS) for 5 min, blocked with defat milk in PBS for 1 h, washed three times with PBS, and incubated with affinity-purified anti-CpeIF4A antibody for 1 h. After three washes with PBS, samples were incubated with goat anti-rabbit-IgG conjugated with Alexa Fluor-488 (1:2,000 dilution) (Thermo Scientific, West Palm Beach, FL, USA) for 1 h, followed by three washes with PBS and counterstaining with DAPI (4′,6-diamidino-2-phenylindole; Sigma-Aldrich; 1.0 μg/mL). Slides were washed with PBS for three times, mounted with an antifade mounting medium (Beyotime, Shanghai, China), sealed with nail polish, and observed under Olympus DP72 research microscopy. Images were saved in TIFF files and processed with Adobe Photoshop. Image signal levels and gamma were properly adjusted to enhance clarity without altering the features and interpretation of the data.

### Effect of Roc-A on the translation and transcription in sporozoites during excystation

#### Effect of Roc-A on the translation of *C. parvum* during excystation

The inhibition of CpeIF4A activity by Roc-A blocks the translational initiation, which would lead to the suppression of protein synthesis in *C. parvum*. This was tested by western blot analysis that quantified the protein contents in excysting sporozoites, for which a sufficient amount of pure parasite materials could be practically obtained. The in vitro excystation procedure was performed by incubating oocysts in RPMI-1640 containing 7.5 mg/mL taurocholic acid at 37 °C for a duration of 2 h in 96-well microplates (10^6^ oocysts in 100 μL per well), in the absence or presence of Roc-A at specified concentrations. To ensure that the reduction of protein contents was truly due to the inhibition of translational initiation, rather than to the death of sporozoites, we first determined that the sporozoites under the above described excystation condition were tolerant to Roc-A treatment up to 100 μM with no or little effect on viability by qRT-PCR detection of Cp18S rRNA (**Fig. 6A**). We also confirmed that Roc-A at 10 μM had no or little effect on the morphology of excysted sporozoites and the distribution of CpeIF4A in the sporozoites (**Fig. 6B**).

We then evaluated the effect of Roc-A at concentrations up to 10 μM on the protein synthesis in excysting sporozoites by western blot analysis. For each treatment, excysted sporozoites (equivalent to 10^6^ oocysts in 50 μL) were centrifuged to collected pellets and supernatants. The supernatants were mixed with sample loading buffer for SDS-PAGE. Pellets were first lysed in RIPA buffer (50 μL) containing protease inhibitor cocktail, followed by incubation on ice for 10 min, vortex for 1 min, and centrifugation to collect supernatants, which were mixed with sample loading buffer. The two preparations, which represented secreted and non-secreted proteins, were subjected to SDS-PAGE (20 μL/lane in 10% gel) and blotted onto nitrocellulose membranes.

The blots were detected using a rabbit antiserum raised against total protein extracted from *C. parvum* oocysts (pan-*Cryptosporidium* antibody). The grayscale intensities of the whole lanes for non-secreted proteins or the specified lane for secreted protein were measured using Fiji ImageJ2 (https://imagej.net/), and plotted against the Roc-A concentrations for nonlinear regression to compute the IC_50_ values.

#### Effect of Roc-A on the transcription of *C. parvum* during excystation

The effect of Roc-A on transcription was studied by transcriptomics. Free sporozoites were prepared by excystation for 1 h, resuspended in RMPI-1640 medium, and divided into six aliquots (2×10^7^ sporozoites per aliquot). Samples were treated with 0.15 μM Roc-A or diluent (0.5% DMSO) at 25 °C for 2 h (N = 3 per treatment group). Sporozoites were collected by centrifugation, resuspended in diethylpyrocarbonate (DEPC)-treated nuclease-free water, snap-frozen in liquid nitrogen, placed in dry ice, and shipped to Shanghai Personalbio Technology (Shanghai, China) for RNA isolation and RNA-seq as briefly described below.

Total RNA was isolated using the TRIzol Reagent (Invitrogen), The quality, integrity and quantity of the isolated RNA were assessed using NanoDrop spectrophotometer (Thermo Scientific). Each sample used 3 μg RNA for constructing cDNA libraries, which included the isolation of mRNA using poly-T-oligonucleotide-attached magnetic beads, fractionation by divalent cation chromatography, synthesis of first strand cDNA using random oligonucleotides, removal of the RNA strand by RNase H, synthesis of second strand DNA, and ligation of Illumina PE adapter oligonucleotides. The preparations were fractionated, from which 400–500 bp fragments were collected for enrichment for those containing adapters on both ends by PCR. Products were purified and quantified using the Agilent high sensitivity DNA assay on a Bioanalyzer 2100 system (Agilent). The cDNA preparations were sequenced on Illumina NovaSeq 6000 system.

Raw reads were filtered to obtain high quality clean reads that were mapped to the *C. parvum* reference genome using HISAT2 (v2.1.0). Expression levels were analyzed by comparing the counts of clean reads by HTSeq (v0.9.1) and converted to FPKM (fragments per kilobase of transcript per million mapped reads). Differentially expressed genes (DEGs) were identified using DESeq (v1.38.3) (cutoffs for statistical significance: |log2(Fold-change) >2.0 and p-value or adjusted p-value (q) <0.05).

## Data analysis and statistics

For all assays, at least two independent experiments were conducted for each experiment condition, except for the animal experiments and transcriptomic analysis that were performed once. Each experimental conditions included three or more biological replicates, and two technical replicates for qRT-PCR. In vitro and biochemical data were analyzed with GraphPad Prism (version 10.0 or higher; Boston, MA, USA) or as specified. Transcriptomes were analyzed using various software packages as specified. Statistical significances were determined by two-way analysis of variance (ANOVA) and appropriate multiple t-tests between group pairs based on the datasets or as specified.

## Ethics Statement

All animal experimentation followed the Guide for the Care and Use of Laboratory Animals of the Ministry of Health, China. The research protocols were reviewed and approved by the Animal Welfare and Research Ethics Committee of Jilin University (AUP No.2020-1Z-20).

## Acknowledgements

We thank Ms. Chenchen Wang at the Institute of Zoonosis, Jilin University for technical assistance in immunostaining related experiments.

## Author Contributions

Conceptualization: Guan Zhu, Meng Li.

Data curation: Meng Li, Dongqiang Wang, Guan Zhu.

Formal analysis: Meng Li, Dongqiang Wang, Guan Zhu.

Funding acquisition: Jigang Yin, Guan Zhu.

Investigation: Meng Li, Dongqiang Wang, Pingwei Li, Beibei Zou

Methodology: Meng Li, Dongqiang Wang, Pingwei Li, Guan Zhu.

Supervision: Guan Zhu, Jigang Yin.

Writing – original draft: Meng Li, Guan Zhu.

Writing – review & editing: Meng Li, Guan Zhu.

## Supporting information

**S1 Table.** Transcript abundance comparison of *CpeIF4A* gene with *CpeIF4A1* and one of the DEAD-box (DDX) domain-containing proteins clustered with DDX19/25 proteins (cgd8_4750).

| Sample | CpeIF4A (cgd1_880) |  | CpeIF4A1 (cgd7_3940) |  | DDX19/25 (cgd8_4750) |  |
| --- | --- | --- | --- | --- | --- | --- |
|  | TPM | Fold | TPM | Fold | TPM | Fold |
| Oocysts- sense - unique | 4,265.2 | 1.0 | 58.0 | -73.5 | 165.5 | -25.8 |
| Pooled in vitro infection samples - unique | 6,239.5 | 1.0 | 125.3 | -49.8 | 355.9 | -17.5 |
| Asexual - unique | 2,422.4 | 1.0 | 42.1 | -57.6 | 114.1 | -21.2 |
| Female in vitro culture - unique | 600.9 | 1.0 | 7.3 | -82.1 | 22.8 | -26.4 |
| Female in vivo mouse - unique | 1,046.5 | 1.0 | 14.3 | -73.3 | 31.6 | -33.1 |
| Sporozoites - unique | 8,893.7 | 1.0 | 150.5 | -59.1 | 233.8 | -38.0 |
| 24 hr culture - unique | 1,028.5 | 1.0 | 32.2 | -31.9 | 112.2 | -9.2 |
| 48 hr culture - unique | 466.3 | 1.0 | 14.6 | -31.9 | 48.3 | -9.7 |

\*Transcriptomic data were extracted from CryptoDB (https://cryptodb.org/). Transcript abundance is expressed as transcripts per million (TPM). Fold changes were calculated using CpeIF4A (cgd1\_880) as the baseline. Positive values indicate fold increases relative to CpeIF4A, while negative values indicate fold decreases.

**S2 Table.** Proteomic abundance comparison of CpeIF4A with CpeIF4A1 and one of the DEAD-box (DDX) domain-containing proteins clustered with DDX19/25 proteins (cgd8_4750).

| Protein name | Total sequences |  | Unique sequences |  | Sum spectra |  |
| --- | --- | --- | --- | --- | --- | --- |
|  | Count | Fold | Count | Fold | Count | Fold |
| CpeIF4A (cgd1_880) | 53 | 1.0 | 26 | 1.0 | 115 | 1.0 |
| CpeIF4A1 (cgd7_3940) | 1 | -53.0 | 1 | -26.0 | 1 | -115.0 |
| DDX19/25 (cgd8_4750) | 9 | -5.9 | 5 | -5.2 | 10 | -11.5 |

\*Proteomic data were sourced from CryptoDB (https://cryptodb.org/). Proteomic abundances are combined mass spectrum counts from various oocyst and sporozoite samples as described in the CryptoDB. Fold changes were calculated using CpeIF4A (cgd1\_880) as the baseline. Positive values indicate fold increases relative to CpeIF4A, while negative values indicate fold decreases.

**S3 Table.** Mouse health scoring scales.

| Parameters | Score |  |  |
| --- | --- | --- | --- |
|  | 0 | 1 | 2 |
| **Fur condition** | **Normal**<br>Smooth and shiny, no frizz. | **Moderate**<br>Slightly frizzy, generally less shiny. | **Severe**<br>Messy and frizzy; has lost luster. |
| **Body weight** | **Normal to 1% weight loss**<br>Daily weight is increased, unchanged, or decreased by <1%. | **Slight to moderate**<br>Daily weight loss between 1% and 10%. | **Severe weight loss**<br>Daily weight loss >10%. |
| **Hunchbackedness** | **Normal**<br>Active movement and elongated posture. | **Slight to moderate**<br>Not particularly active, their backs slightly arched. They may sway from side to side when they walk. | **Abnormal**<br>Severe arching of the back and reluctance to move. |
| **Behavior** | **Normal**<br>Paying attention to caretakers; actively reacting when caretaker's hands enters the cage. | **Mildly to moderately depressed**<br>Paying some attention to caretakers; Slow moving in response to visitor's touch to cages. | **Severely depressed**<br>Paying no attention to caretakers; No or little moving even when being touched. |

### References

Whitehead JC, Hildebrand BA, Sun M, Rockwood MR, Rose RA, Rockwood K, et al. A clinical frailty index in aging mice: comparisons with frailty index data in humans. J Gerontol A Biol Sci Med Sci. 2014;69(6):621-32. Epub 2013/09/21. doi: 10.1093/gerona/glt136. PubMed PMID: 24051346; PubMed Central PMCID: PMCPMC4022099.

Shrum B, Anantha RV, Xu SX, Donnelly M, Haeryfar SM, McCormick JK, et al. A robust scoring system to evaluate sepsis severity in an animal model. BMC Res Notes. 2014;7:233. Epub 2014/04/15. doi: 10.1186/1756-0500-7-233. PubMed PMID: 24725742; PubMed Central PMCID: PMCPMC4022086.

---

**S1 Fig.**
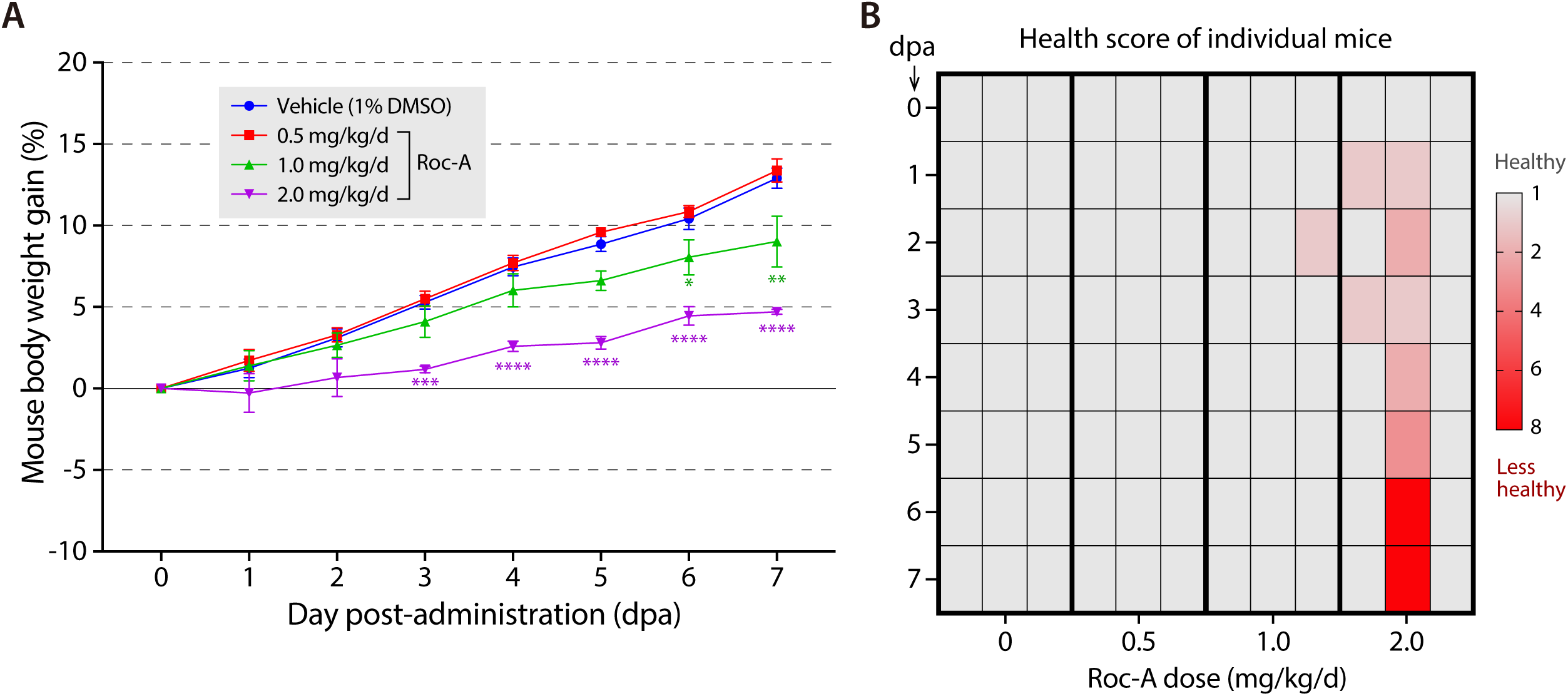
Evaluation of the drug tolerance in mice to Roc-A. C57BL/6 mice (8-wk old; 3 mice/group) were administered by oral gavage with Roc-A at 0 (1% DMSO vehicle), 0.5, 1.0 and 2.0 mg/kg/d in a single daily dose for 7 days. **(A)** Percent daily weight gains in mice from 0 to 7 dpa (day post-administration). Bars indicate standard error of the mean (SEM). * = p <0.05, ** = p <0.01, *** = p <0.001, and **** = p <0.0001 by Tukey’s multiple comparison test (vs. vehicle control). **(B)** Mouse health scores calculated based on the survival, weight gains, fur condition, hunchbackedness, and animal attitude scales (also see Materials and Methods for detail).

**S2 Fig.**
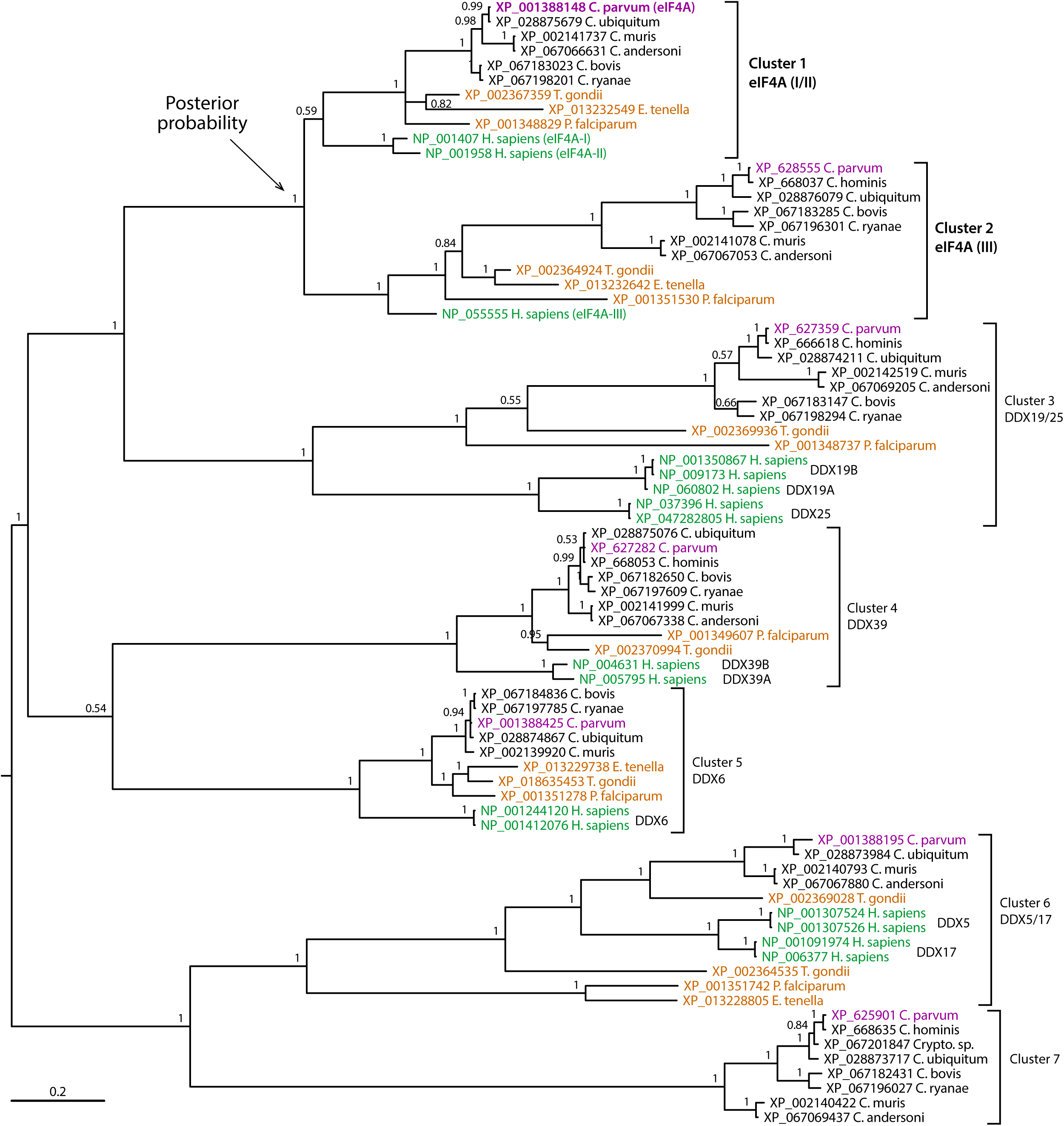
Phylogenetic tree of CpeIF4A and closely related orthologs from various *Cryptosporidium* species, selected apicomplexan species, and humans. The displayed is a consensus tree inferred by Bayesian inference (BI) analysis, which separates the sequences into seven clusters. Six clusters contain human orthologs, including eIF4A-I/II, eIF4A-III, DDX19/25, DDX39, DDX6, and DDX5/17. The seventh cluster appears unique to *Cryptosporidium* (absent in humans and other apicomplexans), but closely related to DDX5/17 subfamily proteins.

**S3 Fig.**
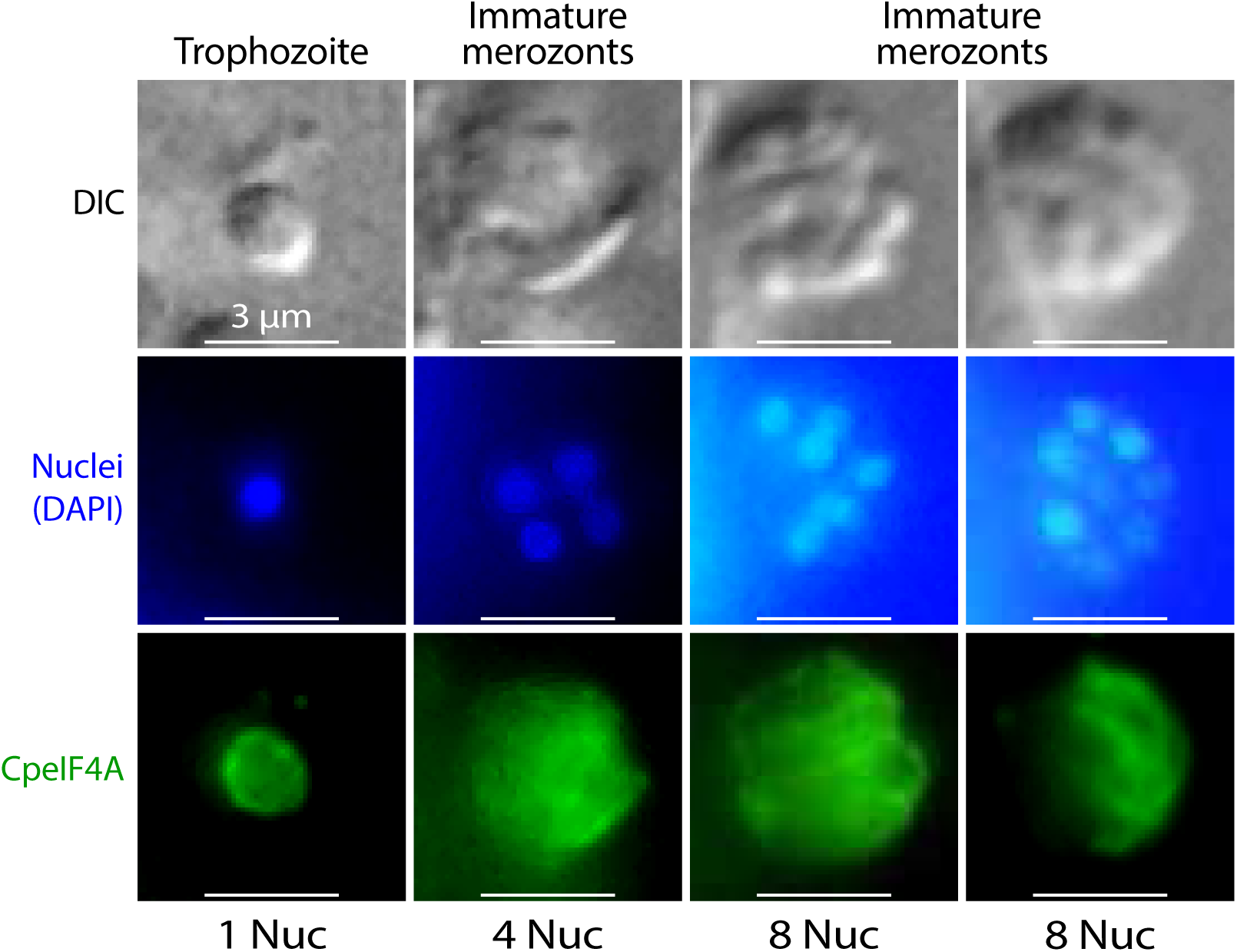
Immunostaining of CpeIF4A in intracellular *C. parvum* cultured with HCT-8 cells. Host cell monolayers infected with *C. parvum* were fixed with paraformaldehyde (4%) and stained with affinity-purified anti-CpeIF4A antibody. Nuclei (Nuc) were counterstained with DAPI. The images show that CpeIF4A is distributed in the cytoplasm with stronger signals under the plasma membrane in varied developmental stages during merogony.

**S4 Fig.**
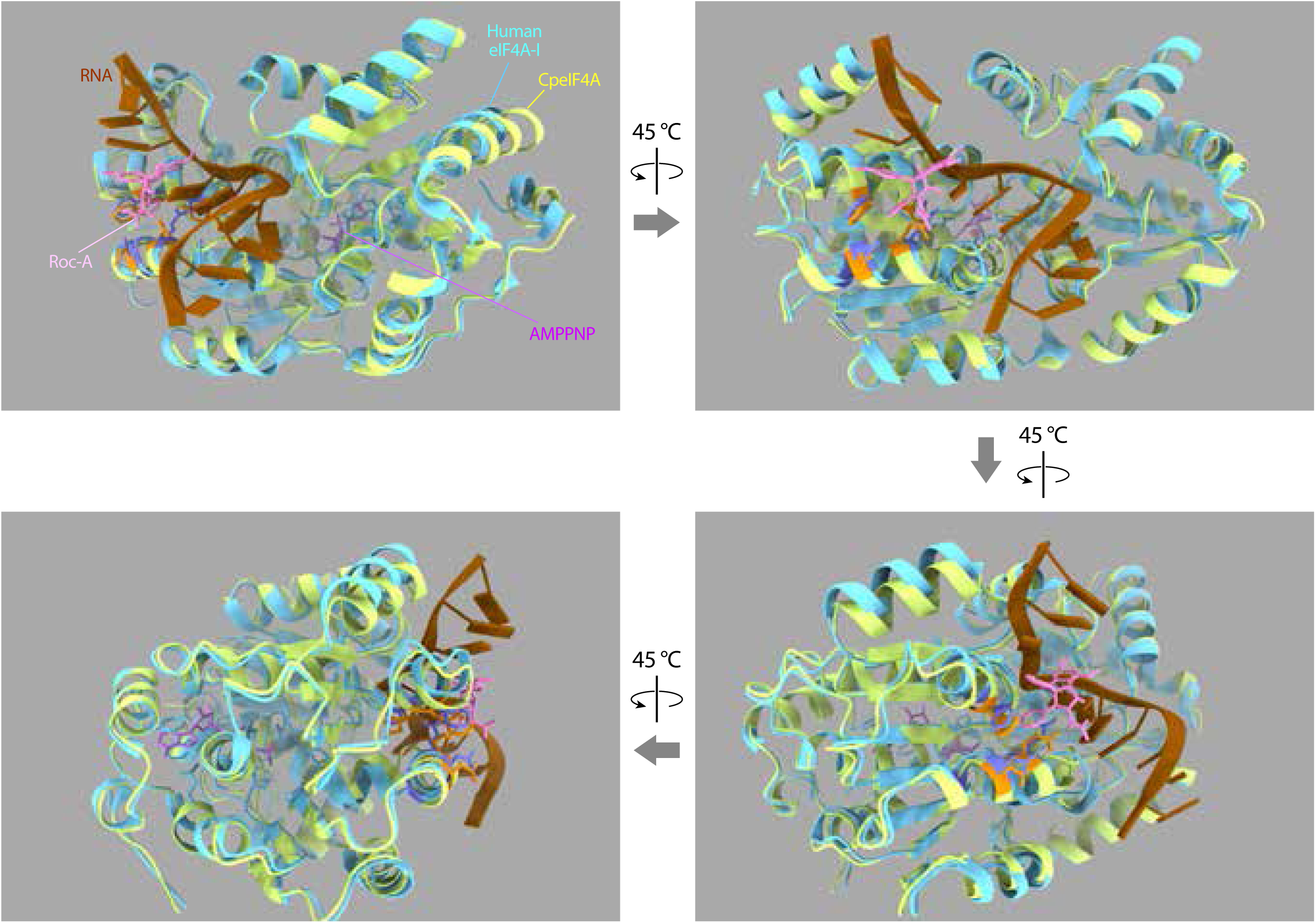
Structure of CpeIF4A superimposed with human eIF4A-I in complex with the ATP analog (AMPPNP), Roc-A and polypurine RNA. CpeIF4A structure is predicted by AlphaFold (AlphaFoldDB entry: A3FQC6) and superimposed with the crystal structure of the human eIF4A1-ATP analog-Roc-A-polypurine RNA complex (PDB entry: 5ZC9) using ChimeraX (v1.8). The superimposed structure shows high structural similarity, but subtle difference, between CpeIF4A and human eIF4A-I, predicting the binding of Roc-A at the “bi-molecular cavity” formed by eIF4A-I and a sharply bent pair of purines in the RNA.

